# PRISM-Seq: An Ultra-sensitive Sequencing Approach For Mapping Lentiviral Integration Sites

**DOI:** 10.64898/2025.12.20.695659

**Authors:** Virender Kumar Pal, Marie Canis, Emily Stone, Nathan L. Board, Klara Lenart, Marcilio Jorge Fumagalli, Colin Kovas, R. Brad Jones, Michel Nussenzweig, Frauke Muecksch, Paul D. Bieniasz, Guinevere Q. Lee

## Abstract

Retroviral integration into host genomes is central to both HIV-1 persistence and the safety and function of lentiviral vectors used in gene and cell therapies. However, existing integration site assays remain limited by sensitivity, input requirements, and analytical complexity, and none have been validated at the single-molecule detection limit. Here, we introduce PRISM-seq, an ultra-sensitive workflow for genome-wide recovery of lentiviral-host junctions, paired with BulkIntSiteR, an open-source, fully automated pipeline for integration site annotation. We show that PRISM-seq accurately identifies proviral insertions across diverse genomic contexts, including euchromatin, heterochromatin, and repeat-rich centromeric regions, and detects high-confidence integration events down to a single input template molecule. By systematically characterizing assay- and amplification-associated noise, we developed a five-step quality control framework that removes PCR- and sequencing-derived artifacts. PRISM-seq also enables quantitative clonal tracking through replicate-based sampling and achieves performance comparable to or exceeding high-input assays at substantially reduced cost.

## Introduction

Integration of retroviral DNA into host DNA underlies both HIV-1 pathogenesis and the use of lentiviral vectors in gene therapy. In HIV-1 infection, the site of viral integration predicts viral transcription and reactivation potential^1–3^, while in lentiviral vector-based gene therapies, vector insertional profiles inform safety and clonal dynamics^4–10^.

Existing proviral integration site sequencing assays and bioinformatics workflows (e.g., INSPIIRED^11,12^, VISPA/VISPA2^13^, QuickMap^14^, VISA^15^, MAVRIC^16^, GeIST^17^, IS-Seq^18^, VSeq-Toolkit^19^ face three practical barriers. First, many require relatively high template input, typically ∼1-10 µg of genomic DNA (gDNA) per library^20,21^, are incompatible with single cell sorting, and none were validated down to the single-molecule detection limit nor across distinct genome contexts, including difficult-to-map regions. Second, computational workflows are fragmented, platform-specific, and/or require bioinformatics expertise for read processing and annotation^12,20^. Third, per-sample per-replicate costs remain high due to multi-step library construction and sequencing depth requirements.

Here, we present proviral integration site mapping sequencing (PRISM-seq), an ultra-sensitive lentiviral integration site sequencing assay compatible with both single-copy and bulk-input samples, together with BulkIntSiteR, a fully automated and platform-agnostic bioinformatics analysis pipeline that converts raw sequencing data files into annotated proviral integration sites with a single click.

## Results

### PRISM-seq design

PRISM-seq detects single-molecule inputs and single-copy species in bulk samples. The core of the PRISM-seq design that enabled ultra-sensitive single-molecule compatible detection is bacteriophage phi29-mediated multiple displacement amplification (MDA), which is not commonly used in integration site identification (Fig. 1a and see methods for the origin and foundation assays for PRISM-seq). In this study, reactions were performed using 50-template inputs as a defined starting point for systematic assay characterization; however, the workflow can accommodate substantially higher input levels (∼10,000 proviral templates pre-MDA^22^). MDA-amplified DNA is processed by ligation-mediated PCR (LM-PCR) involving restriction digestion, adaptor ligation, and amplification of both 5′ and 3′ virus-host junctions using virus-and adaptor-specific primer pairs (Fig. 1a, Fig. S1 and S2a). The PRISM-seq primer sets were designed for broad compatibility across lentiviruses, including HIV and lentiviral vector delivery systems (Fig. S2b)^23–27^. PRISM-seq amplicons containing viral-host junctions can be sequenced on any next-generation sequencing (NGS) platform, and the reads obtained were processed using our open-source computational pipeline, BulkIntSiteR, which was developed for the automated mapping and annotation of integration sites (Fig. 1b).

**Figure 1.**
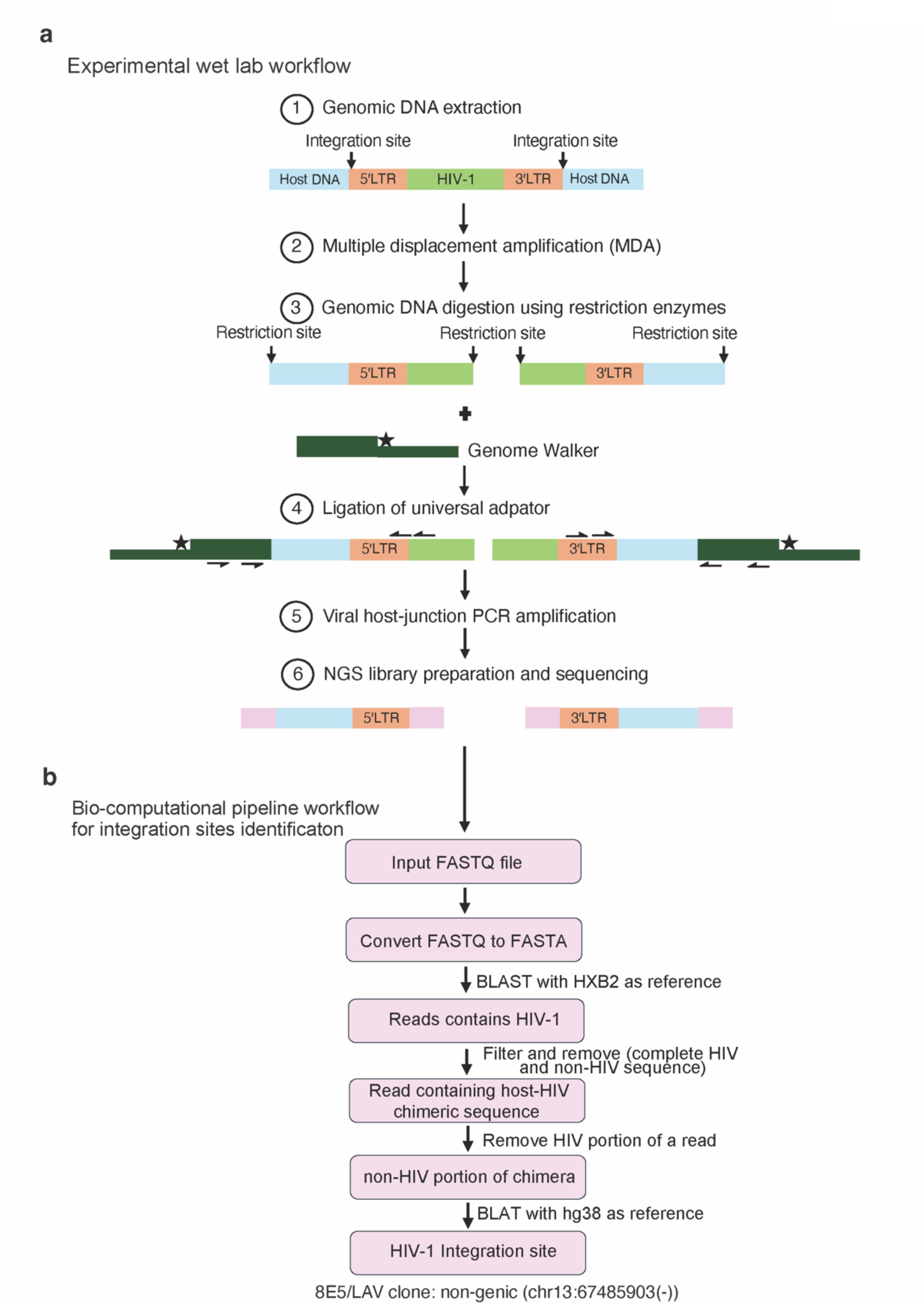
Experimental and bioinformatic workflow for proviral integration sites identification using PRISM-seq. **(a)** Experimental workflow of PRISM-seq. **(b)** Bioinformatics steps implemented in the BulkIntSiteR pipeline to process raw sequencing data and generate annotated proviral integration site outputs. *NGS, next-generation sequencing*.

### Identification of proviral integration sites across the diverse human genome landscape

We first benchmarked PRISM-seq/BulkIntSiteR using the 8E5/LAV cell line, which harbors a single HIV-1 provirus at a known intergenic locus^28^. BulkIntSiteR normalizes junction coordinates to the 5′ LTR convention, accounting for the characteristic 4-bp offset between the 5′ and 3′ LTR junctions^29^. Using PRISM-seq, we identified a single viral integration site in 8E5/LAV cells on chromosome 13 at position 67,485,907 in the negative strand, consistent with published data^28^. We did not observe any low-abundance secondary integration sites (noise).

The single provirus in 8E5/LAV cells is associated with heterochromatin (Supplementary Table 1). However, viral integration can occur in diverse regions of the human genome, including actively transcribing genes^30^, and in repeat-rich, telomeric, and centromeric satellite regions^3,6,31,32^. We generated 14 putative singly-infected Jurkat cell clones (Clones #1-14) using low-MOI (0.3) infection with a replication-deficient HIV-1 reporter (V1/SBP-GFP), followed by sorting of GFP/SBP-positive cells, limiting dilution, and expansion via culturing. Proviral integration sites in these 14 putative clones were expected to be mostly in genes since HIV-1 integration favors actively transcribing genomic regions^30^. To evaluate PRISM-seq performance within centromeric regions typically associated with difficult-to-map satellite DNA repeats, we analyzed two additional HIV-1-infected primary cell clones (#15 and #16) previously found to each contain a single provirus integrated within centromeric regions^32^.

Without noise removal, our raw data showed 1-70 (min-max) unique proviral insertion sites in clones #1-16 (Fig. S3a-c and S4a-c). As each putative clone is expected to harbor a single proviral integration site, the detection of multiple unique integration sites per clone may indicate assay-associated noise, which is consistent with the presence of noise reported in other integration site sequencing assays^20^. We therefore used this clonal sample set as an internal benchmark to systematically evaluate assay noise.

An initial quality-control filter designated each of our raw integration sites as high- or low-confidence. High-confidence was defined if an integration site (i) contained precise viral DNA 5′ or 3′ ends linked to a human sequence without gaps/missing bases, and (ii) was detected by both 5′ LTR and 3′ LTR viral-host junctions or (iii) had greater than 10 reads when identified only by either 5′ LTR or 3′ LTR. All other integration sites were designated “low-confidence”.

Among clones #1-14, 9/14 (64%) yielded a single high-confidence integration site (Fig. S3c). Two of the 14 clones, #6 and #10, showed multiple high-confidence integration sites, consistent with expectations from Poisson statistics for infections at MOI 0.3^33^. Clones #8, #11, and #12 had more than one high-confidence integration sites, and these integrations were found in the same gene for the respective clone, which we explored in-depth in the next section for potential assay-associated artifacts. We used two human reference genomes: hg38 and T2T-CHM13^34^, and their results were 100% concordant for clones #1-14. For clones 15 and 16, our data showed multiple high-confidence integration sites with proviruses integrated within the centromeric repeat regions of chromosomes 9 (8 sites) and 22 (2 sites), respectively (Fig. S4c), which were only detected using T2T-CHM13 as the reference genome, while hg38 returned no maps. Due to the clonal nature of these samples, we suspected repeat-region-driven ambiguity in mapping. All ambiguities were further explored in the next section.

Across all 16 analyzed clones, high-confidence proviral integration sites were distributed within 12 chromosomes spanning genic, intergenic, and repeat-rich centromeric satellite regions, confirmed by the chromHMM model^35^ based on distinct histone-mark ChIP-seq datasets from the ENCODE database^36^ (Supplementary Table 1). The average distance between proviral integration sites and the nearest accessible chromatin region (ATAC-seq peak) varied widely, ranging from 292 bp to 278,541 bp (Supplementary Table 1). These findings reflect the diverse genomic landscape of proviral integration recovered by PRISM-seq.

### Implementation of additional quality control filtering criteria

In 15 of 16 analyzed clones, we detected 174 low-confidence secondary integration sites in addition to the described high-confidence integration sites. We next characterized this putative noise. Of these low-confidence integration sites, 57/174 (33%) mapped to the same gene or within ±10 kb relative to the corresponding high-confidence integration sites (Fig. S3a-c, S4a-c, S5a-b, and Supplementary Table 2). The remaining low-confidence sites were either in intergenic regions or in distinct genes. Given the clonal nature of most of the samples, these sites may constitute artifacts. All junction sequences were subjected to manual examination as described below.

To further investigate these potential artificats, we performed manual alignment of the host sequences of low-confidence integration sites, and showed that many differed from the corresponding high-confidence site of the clone only by short indels or 1-10 bp shifts in the junction sequence (e.g., clones 1, 2, 4, 8, 9, 10, 11, 13, 15, and 16) (Fig. S6-S12). These cases are likely artifact signatures previously highlighted to be found in integration site sequencing assays, caused mainly by PCR recombination, whole-genome amplification artifacts, sequencing errors, or ambiguous mapping in repeat-rich chromosomal regions^20,37,38^. Collectively, the short indel cases, same gene integration in a putative clone, and low read-count (≤10) when captured by only one proviral end accounted for 174/174 (100%) of the low-confidence integration sites detected among clones #1-16 (Supplementary Table 2). Together, these observations demonstrated that low-confidence integration sites arose from predictable, assay-associated artifacts, providing a rational basis for systematic quality-control filtering.

We therefore implemented a five-step quality control filter strategy to improve the specificity of PRISM-seq (Fig. S13). Steps 1-4 are fully automated, and the accompanying script is included with the BulkIntSiteR package; Step 5 requires manual curation. In this study, steps 1-4 removed 94% (164/174) and step 5 removed an additional 6% (10/174) of the noise in our data. Step 5 is especially critical when proviruses are integrated within repeat-rich genomic regions: without it, 7 of 8 (88%) and 1 of 2 (50%) integration sites were called as high-confidence sites for the two clones (#15 and #16) with centromeric proviral insertion site (Fig. S14a-e). Incorporating step 5 yielded a single high-confidence integration site for each clone. All integration sites that pass the final filter are designated as “high confidence integration sites” and are used for downstream analysis.

### PRISM-seq detection efficiency and cumulative recovery in samples containing multiple integration sites

Next, we examined whether PRISM-seq can accurately detect integration sites in bulk input samples with more than one unique integration site, each with varying abundance. Variation in template input copies, as well as multiple authentic proviral insertions, can limit detection efficiency and accuracy due to PCR competition and recombination^12,20,39^. To model this scenario, we generated two artificial gDNA pools composed of clones #1-14 (Pool 1 and Pool 2), each with integration sites represented at different relative input frequencies, and used them to assess the detection efficiency of PRISM-seq in a mixed template context (Fig. 2a-b).

**Figure 2.**
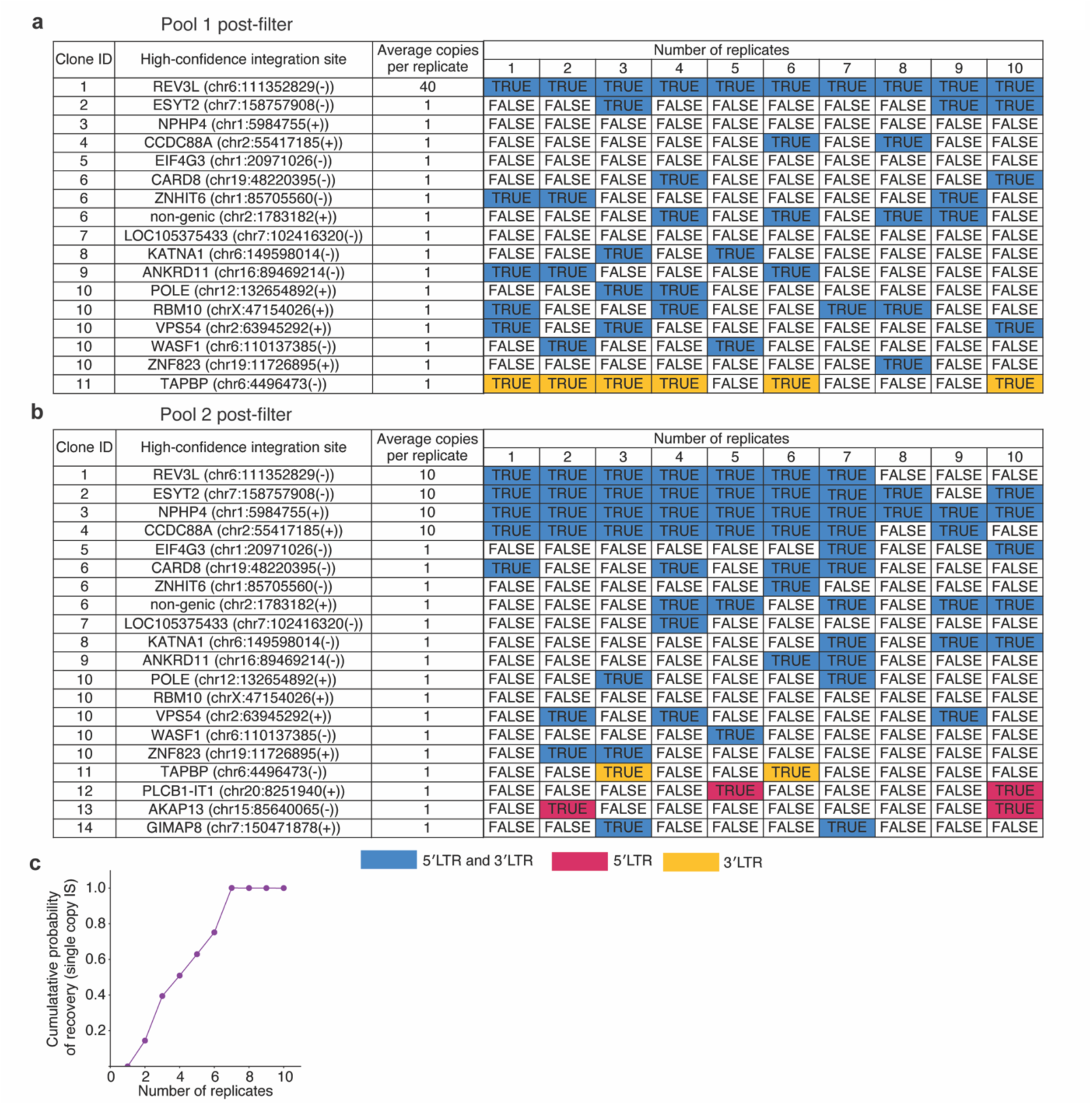
Detection efficiency of PRISM-seq in artificial mixes. **(a and b)** Two artificial pools were constructed to test PRISM-seq’s ability to recover integration sites in samples with multiple unique integration sites at varying relative abundance. Pool 1 contained 11 of the 14 clones #1-14, whereas pool 2 contained all 14 clones #1-14. Each pool was divided into ten independent genomic DNA sampling replicates and independently subjected to PRIMS-seq. The “average copies per replicate” column shows the copy number of high-confidence integration sites present in the artificial pool 1 (a) and pool 2 (b), respectively. The colored matrix shows the recovery of expected high-confidence integration sites across 10 replicates. The color highlights if a high-confidence integration site was recovered by both 5′LTR and 3′LTR (blue), or exclusively by either 5′LTR (red), or 3′LTR (yellow) viral-host junction. True/false denotes if a high-confidence integration site was recovered by either proviral end. **(c)** Cumulative recovery of single-copy clone integration sites after adjusting for Poisson sampling expectation across 10 replicates for pool 2. Clones #6 and #10 were associated with more than one integration site and were excluded from this analysis. *IS, integration sites*.

We first estimated the number of molecules containing viral integration sites for each clone based on gDNA input mass, assuming a clonal population structure and using an established approximation of ∼150,000 diploid human cells per microgram of genomic DNA^40^. Using these estimates, we constructed two master pools: Pool 1 consisted of ∼400 copies of clone #1 combined with ten additional clones, each represented at ∼10 copies, whereas Pool 2 consisted of ∼100 copies each of clones #1-4 combined with ten additional clones, each represented at ∼10 copies. These two mixes were each distributed into 10 replicate reactions. Therefore, effectively, Pool 1 contained clone #1 at an average of ∼40 copies per reaction, while clones #2-11 were present at approximately single-copy input per reaction, whereas Pool 2 contained clones #1-4 at ∼10 copies per reaction and clones #5-14 at approximately single-copy input per reaction. The automated quality-control filter steps 1-4 outlined in the previous section were applied to all results. Note that this experimental design represents a limiting-dilution strategy, such that the actual number of template molecules distributed into each reaction followed Poisson statistics, with mean (λ, lamda) values of 40, 10, or 1 copy per reaction.

With this 10-replicate setup, the abundant 40- and 10-copy integration sites were detected with 100% efficiency in both pools (Fig. 2a-b and S15-17). As for single-copy integration site templates, each replicate reaction contained an average of one HIV template. Under a Poisson sampling model, the probability that a given well received at least one template molecule is 63%; however, when multiple replicate reactions were performed, the cumulative probability of observing at least one positive well increased with the number of replicates. Therefore, detection efficiency for single-copy integration sites was evaluated in the context of replicate-level cumulative recovery rather than single-well sampling probability. After excluding clones with multiple integration sites (clones #6 and #10), three single-copy integration sites (clones #3, #5, and #7) were not detected in Pool 1 across all replicate reactions (Fig. 2a). Under a Poisson sampling model, assuming an average of 10 template copies per 10 cumulative reactions (λ = 10), the probability of observing zero molecules in a given set of 10-replicate is <0.1%, corresponding to a >99.9% probability that at least one molecule is present. This indicates that stochastic loss at the level of individual reactions is unlikely to explain the observed lack of signal in Pool 1. Therefore, the absence of these clones in Pool 1 is most consistent with either (i) their under-representation or absence during pool construction, or (ii) clone-specific amplification or detection failure. In contrast, all three clones #3, #5, and #7 were also input as a single template per reaction in Pool 2 but were robustly detected (Fig. 2b), ruling out integration-site-specific amplification or detection limitations and strongly implicating upstream sampling during Pool 1 preparation as the primary cause of their loss. Consequently, detection efficiency calculations for rare (single-copy) integration sites were based on Pool 2 data. Using replicate-level cumulative recovery and correcting for Poisson sampling expectations, 75% of single-copy integration sites were detected by six replicates, with complete (100%) recovery achieved by seven replicates, despite the presence of abundant competitor templates (Fig. 2c). Overall, these findings demonstrate that PRISM-seq can reliably recover integration sites in a sample expected to contain both rare and abundant templates.

### Estimation of the extent of clonal expansion using PRISM-seq

Repeated detection of the same integration site across pre-MDA replicates signifies that the original sample contains multiple copies of a given viral-host junction (Fig. 3a). In a clinical context, such an observation would indicate clonal expansion of infected cells. We reasoned that the frequency with which an integration site appears across PRISM-seq replicates would be an indirect quantitative measure of the relative size of the clone harboring that integration.

**Figure 3.**
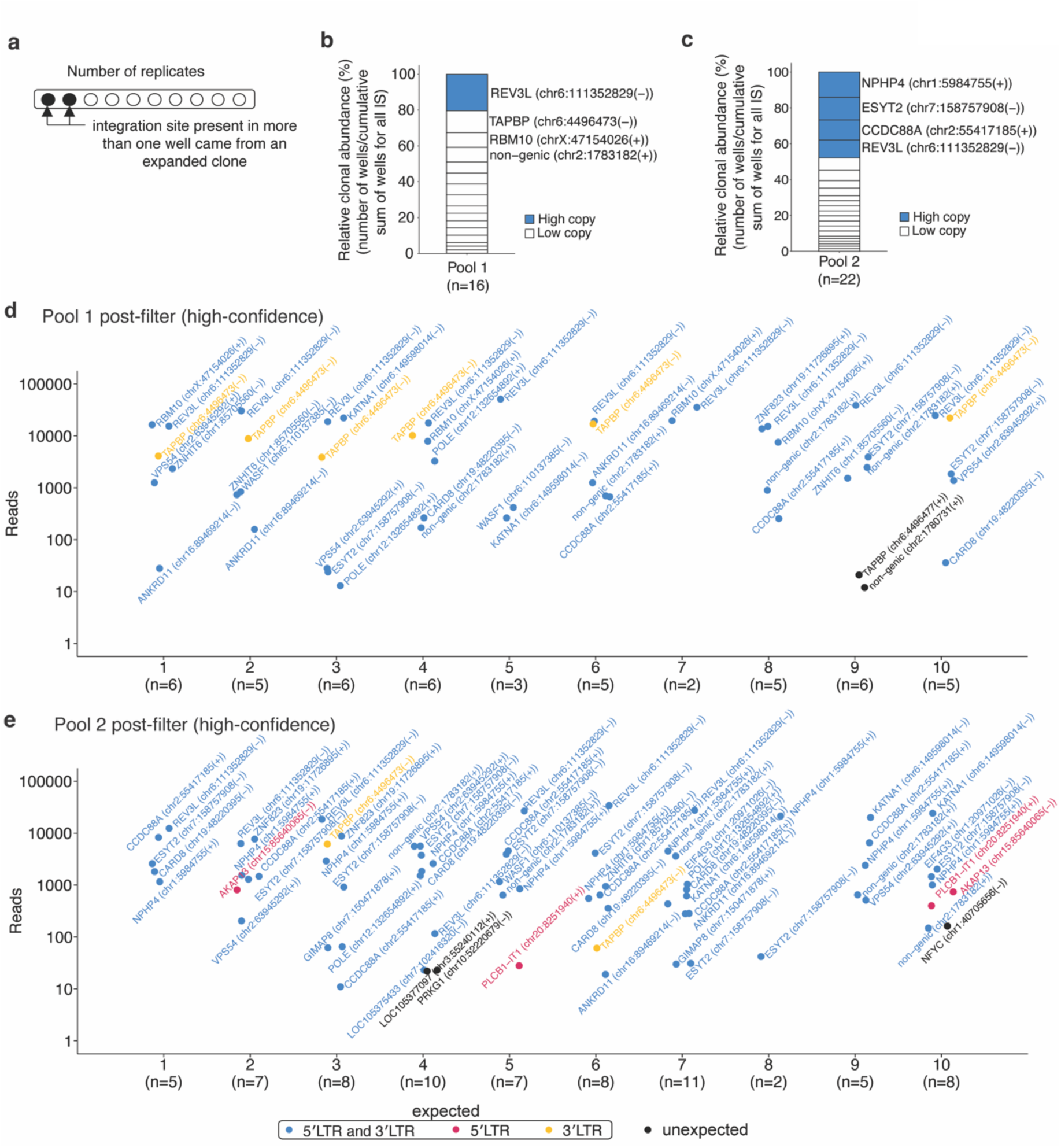
Post-quality filter recovery of high-confidence integration sites and estimation of clonal abundance. **(a)** Schematics showing the identification of expanded clones using PRISM-seq. We used the post-filtered data to evaluate integration site authenticity and to estimate clonal abundance in pool 1 and pool 2, containing multiple integration sites. **(b and c)** Post-filter estimation of the relative clonal abundance of individual clones found in pool 1 and pool 2. After applying steps1-4 of the quality-control filter, we recovered a total of 16 and 22 high-confidence integration sites in pool 1 and pool 2, closely matching the expected input sites (17 for pool 1 and 20 for pool 2), with false-positive calls limited to 2 of 16 (12.5% noise) and 3 of 22 (13.6% noise), respectively. The top 4 abundant clones are labelled in the stack bar plot. **(d and e)** The dot plots show the total number of integration sites retrieved from individual sequencing replicates of pool 1 (a) and pool 2 (b), respectively. Expected high-confidence integration sites are highlighted if recovered by both 5′LTR and 3′LTR (blue), or exclusively by either 5′LTR (red), or 3′LTR (yellow) viral-host junction. Unexpected integration sites are colored in black. *IS, integration sites*.

As proof of principle, we applied this framework to the post-filtered datasets for pools containing multiple integration sites. For each integration site, we counted the number of replicate reactions in which it was recovered. Integration sites detected in multiple replicates are highly likely to represent true biological events, whereas those observed only once are more likely to reflect residual noise. As anticipated, the dominant clones, present at ∼400 copies in Pool 1 and ∼100 copies in Pool 2, were detected in the largest number of replicate reactions and therefore exhibited the highest inferred clonal abundance (Fig. 3b-3e). In Pool 1, the high-copy integration site was recovered in all 10 replicate wells, corresponding to 20% of the total relative clonal abundance, whereas lower-copy integration sites were detected in fewer wells (1-6 wells; 2-12% clonal abundance) (Fig. 3b and 3d). Similarly, in Pool 2, high-copy integration sites were recovered in 7-10 of 10 replicate wells, accounting for 10-14% of the total relative clonal abundance, while low-copy integration sites were detected in fewer replicates (1-5 wells; 1-7% clonal abundance) (Fig. 3c and 3e).

### Application to longitudinal clinical samples from a person with HIV-1 on ART

To demonstrate the clinical utility of PRISM-seq, we longitudinally sampled CD4^+^ T cells from a study participant living with HIV-1 before antiretroviral therapy initiation (pre-ART), and 3 months, 1 year, and 7 years post-ART (Supplementary Table 3). Each sample was expected to contain multiple unique integration sites originating from infected cells exhibiting variable levels of clonal expansion^41–44^. Using 2-5 replicates each containing 50 proviral copies, and thus only sampling 100-250 HIV-containing templates per time point, we recovered 69, 67, 11, and 17 high-confidence integration sites from each time points, respectively (Fig. 4a). The proportion of proviruses integrated into genic or intergenic regions remained stable over time (Fig. 4b). The fraction of proviruses integrated into ZNF genes increased from 2%-3% (pre-ART and 3 months) to 9% (1 year) and 18% (7 years) post-ART time points (Fig. 4c), consistent with previous reports indicating that proviruses persisting in ZNF genes and transcriptionally repressive genomic contexts are more likely to be transcriptionally silent and capable of long-term persistence^6,31^.

**Figure 4.**
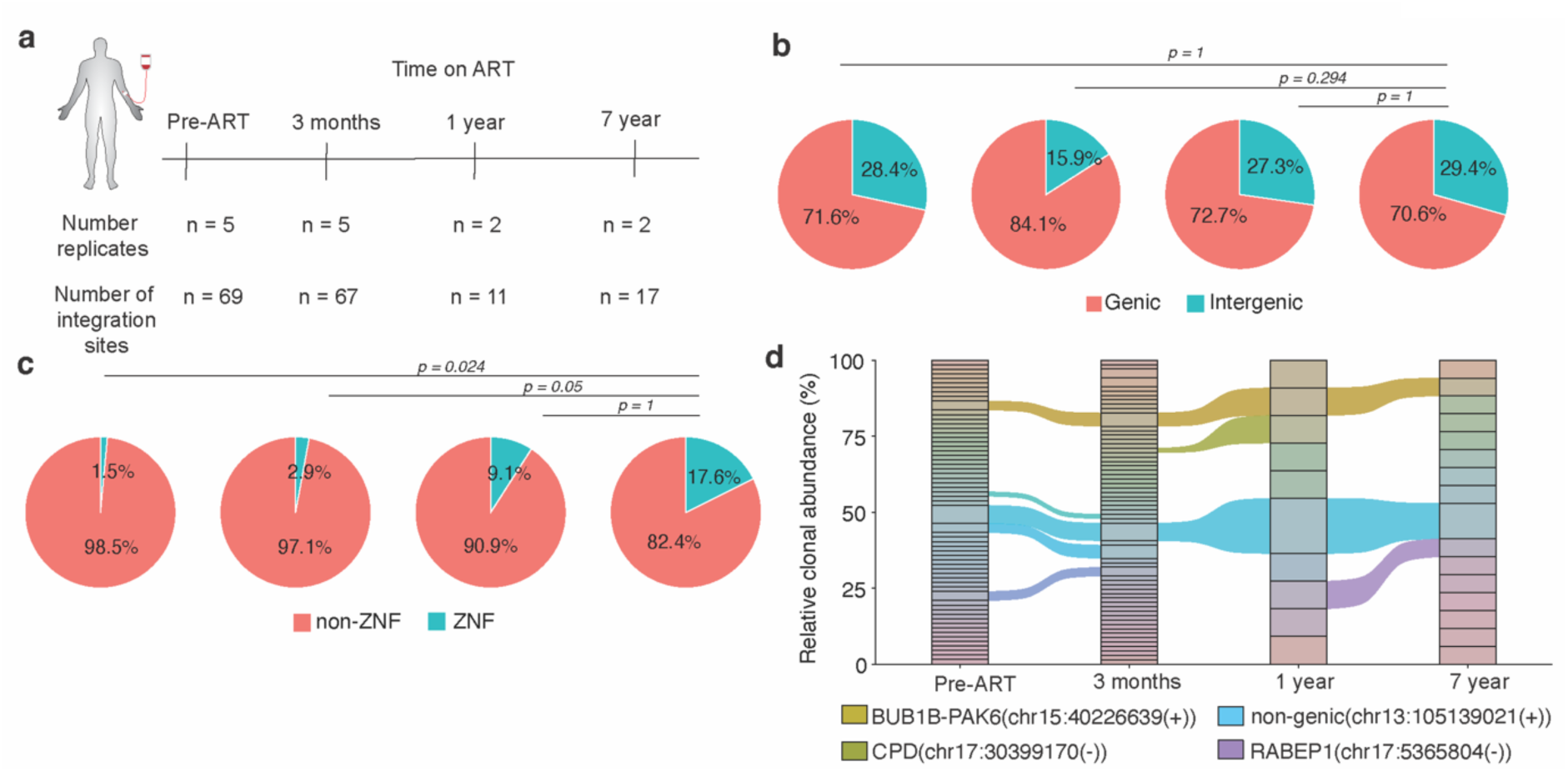
Longitudinal clonal tracking and identification of proviral integration sites in a person living with HIV-1. **(a)** Number of unique integration sites identified at each time point using PRISM-seq from a person living with HIV-1 under long-term antiretroviral therapy (ART) treatment. Across the number of replicates performed, we sampled in total 250, 250, 100, and 100 HIV-containing templates at each indicated time point. **(b)** Precent genic and intergenic distribution of proviral integration sites. **(c)** Distribution of proviruses in ZNF and non-ZNF genes in the genome. P-value denotes Fisher’s exact test. **(d)** Clonal tracking of HIV-1 infected cell clones over time under ART. Shared clones identified at all the time points are labelled with their respective proviral integration sites.

Considering that repeated detection of the same integration site in multiple replicates indicates a higher chance of clonal expansion of the respective cell clone, we used this metric across the longitudinal samples to track persistent clones: we observed identical viral integration sites in BUB1B-PAK6(chr15:40226639(+)) and intergenic region (chr13:105139021(+)) across all 4 time points sampled over 7 years (Fig. 4d), an indicator that HIV-1 infected cells with provirus integrations at these loci were expanded *in vivo* and persisted during treatment over time in this study participant. Overall, our findings show that PRISM-seq could be directly applied to clinical samples that had an extremely low target concentration, and provided a wealth of integration site data and biological insights with minimal sample input.

## Discussion

In this study, we present PRISM-seq and its accompanying software BulkIntSiteR. The assay recovers integration sites from single-molecule input, captures diverse genomic contexts, quantifies clonal expansion, and is compatible with *ex vivo* clinical samples.

Existing bulk integration site assays, including assays INSPIIRED^11,12^, VISPA/VISPA2^13^, QuickMap^14^, VISA^15^, MAVRIC^16^, GeIST^17^, IS-Seq^18^, VSeq-Toolkit^19^, were not validated for single-molecule detection and/or broad performance across challenging genomic contexts. PRISM-seq overcomes these limitations through its MDA-enabled single-molecule capture workflow, which efficiently recovers rare viral-host junctions from minimal DNA input. Although MDA has been used previously in MIP-seq^5^ that approach requires limiting dilution, it is costly, and is not designed for bulk integration site analysis. PRISM-seq provides an economic, bulk-input compatible, sensitive, and scalable platform for high-confidence integration site profiling.

Another key advance of PRISM-seq is the systematic characterization and mitigation of assay-associated noise, an aspect that has not been quantitatively defined in prior proviral integration site sequencing methods^12,20^ nor formalized into a standardized filtering framework. Across biologically pure samples with a single provirus integration site, we identified recurring artifact classes arising from whole-genome amplification, PCR recombination, sequencing errors, and mapping ambiguity in repeat-rich regions. These artifacts manifested as displaced junctions, small indels, low-read viral-host chimeras detected by only one LTR end, and clusters of look-alike integration sites mapping within the same gene or genomic window as the true high-confidence site. Using clonal standards, including centromere-integrated proviruses resolvable only using T2T-CHM13 as a reference genome, we were able to distinguish genuine proviral insertions from amplification- or sequencing-derived noise and develop a five-step quality control pipeline that removes >85-90% of raw artifacts while preserving bona fide integration sites.

Application of PRISM-seq to longitudinal samples from a study participant with HIV-1 revealed persistent clones integrated within transcriptionally repressive, ZNF-rich genomic regions. These findings reinforce prior observations that proviruses can integrate and persist in epigenetically repressed environments, including KRAB-ZNF gene clusters and other heterochromatic regions^2,6,31,45^. The compatibility of PRISM-seq with low sample input and its ability to capture low-abundance, genomically diverse proviruses directly from clinically-derived material highlights its utility for high-resolution reservoir characterization. Furthermore, our companion study^46^ applied PRISM-seq to flow-cytometry-sorted cells derived from a novel mouse model of HIV latency, demonstrating both that proviral integration site is a deterministic factor in proviral latency fate decisions in the absence of external immunological or anti-viral drug selection pressure and that PRISM-seq is compatible with post-sort samples.

PRISM-seq has several limitations. First, clonal expansion estimates currently require replicate reactions and yield relative, rather than absolute, quantification. Second, increasing input concentration beyond ≥50 templates per reaction is possible^22^ but means users should turn off the same-gene filter, as the frequency of true same-gene integrations within a replicate will be higher with increasing template concentrations (step 3 in Fig. S9 and see Monte Carlo simulation Fig S5b). This will result in a modest compromise to specificity. Third, while our quality-filtering framework effectively reduces artifacts, it may occasionally exclude rare authentic sites detected by only one proviral end with low read counts - manual check is always recommended. Finally, PRISM-seq provides precise integration sites mapping but does not assess proviral genome intactness, which remains specific to HIV-1 reservoir studies and should be complemented with assays that evaluate proviral sequence integrity^5,47,48^.

Overall, we have shown that PRISM-seq is an ultra-sensitive, streamlined, and economical method for detecting lentiviral integration sites with single-template resolution. Coupled with its automated bioinformatics pipeline BulkIntSiteR and noise-mitigating steps, the assay delivers a fully integrated workflow for high-confidence integration site identification, filtering, and annotation, especially for rare template detection.

## Method

### Viral-host junction amplification and sequencing

Genomic DNA was extracted (QIAGEN DNeasy kit), quantified for HIV-1 *gag* by droplet digital PCR (Bio-Rad), subjected to whole-genome amplification using multiple displacement amplification (MDA; QIAGEN REPLI-g Single Cell Kit) at 50 copies per reaction, and processed using an adaptor-ligation PCR workflow adapted from the Clontech Lenti-X Integration Site Analysis Kit^5^. Final nested PCR was used to selectively enrich viral-host junction products and sequenced on an Illumina MiSeq platform (2 × 150 bp). Full protocol details, origin of PRISM-seq, reagent specifications, and cost are provided in the Extended Methods. Technical replicates were defined as independent sampling of each DNA extraction (pre-MDA); 10 replicates were used for pooling experiments, 2-5 for clinical samples, and none for 8E5/LAV or clones #1-16.

### Identification of HIV-1 integration site using BulkIntSiteR

Illumina sequencing-derived FASTQ files were processed using BulkIntSiteR, a one-click R-based, fully automated pipeline designed to detect, map, and annotate HIV-1 integration sites in the human genome. BulkIntSiteR is cross-platform and is not restricted to Illumina data. All scripts are available freely on GitHub (https://github.com/guineverelee/BulkIntSiteR/). Briefly, each small read was independently assessed for the presence of high-quality viral ends (containing HXB2 1 or 9719 using BLAST+ suite^49^). For each read positively identified to contain a viral end, the non-viral portion of the read was mapped to the human genome using BLAT^50^. Additionally, we observed a large disparity in junction-containing reads between the 5′ and 3′ LTR integration sites libraries (e.g., for the 8E5/LAV benchmarking experiment), likely reflecting the differences in amplicon composition, relative to Illumina tagmentation and read length. The 5′ LTR amplicon contains ∼681 nt of viral sequence, whereas the 3′ LTR amplicon contains only ∼34 nt. Because tagmentation generates fragments of ∼300 bp on average and the sequencing read length was 150 bp, most 5′ LTR reads fall entirely within the viral portion of the amplicon and never reach the host sequence, which would markedly reduce the number of recoverable viral-host chimeras. See the GitHub BulkIntSiteR documentation for the full algorithm.

### Generation of 14 putative clones with a single integrated provirus

To generate the V1/SBP-GFP HIV-1 reporter virus, 7.5 × 10⁷ 293T cells (ATCC #CRL-3216; RRID: CVCL_0063) were seeded in 875 cm² 5-layer flasks 1 day prior to transfection. 293T cells were transfected with 75 µg pV1/SBP-GFP, 75 µg pCRV1/NLGagPol, and 15 µg pVSV-G (Yee et al., 1994) using polyethyleneimine (PEI). After 12 hours, the medium was replaced with fresh growth medium. Supernatants were collected 48 hours post-transfection, filtered, and concentrated by ultracentrifugation over a 20% sucrose cushion. The pellet containing the virus was resuspended in PBS + 10% BSA, aliquoted, and stored at -80 °C. The viral stock titre (IU/ml) was determined by infecting MT4-LTR-GFP cells and using flow cytometry to assess GFP expression.

For infection, Jurkat and MOLT-4 cells were infected with V1/SBP-GFP^46^ at a multiplicity of infection (MOI) of 0.3. Infected cells were enriched using streptavidin bead-based magnetic selection. Single-cell clones of infected Jurkat and MOLT-4 cells were isolated by limiting dilution by diluting to 0.5 cells per well in RPMI growth medium, and 100 µL of the suspension was seeded into each well of 96-well plates. The clonal outgrowth was monitored microscopically, and wells containing single-cell-derived colonies were transferred to fresh wells for expansion.

### Clinical samples

The study participant with HIV-1 was recruited at the Maple Leaf Clinic in Toronto, Canada. PBMC samples were used according to protocols approved by the respective Institutional Review Boards. CD4^+^ T cells were enriched from total PBMCs using a CD4^+^ T Cell Isolation Kit (STEMCELL Technologies, catalog #17952). Clinical and demographic characteristics of the study participant are summarized in Supplemental Table 3.

### Extended methods (Supplementary)

#### Cost and origin

PRISM-seq is an evolution of our previously published matched integration site and proviral sequencing (MIP-seq) protocol^5^, which itself was adapted from the Clontech LentiX integration site analysis method (catalog #631263). MIP-seq was designed to co-capture HIV integration sites together with full-length proviral genomes at single-input-template resolution. In contrast, PRISM-seq is optimized for both single and multiple-template input and focuses specifically on capturing viral integration sites, enabling substantially higher throughput at dramatically reduced cost. At the time of writing, PRISM-seq offers an economical solution at $280 per sample with detection sensitivity down to a single template and supports up to ∼10000 pre-MDA template input^22^, making the assay broadly accessible. In addition, PRISM-seq is fully supported by a publicly available bioinformatics (BulkIntSiteR) and quality control pipeline, which automates integration site calling, annotation, and quality assessment.

### Input template quantification by Droplet digital PCR (ddPCR)

DdPCR quantifications were performed to estimate the HIV-1 *gag* and *env*-containing proviral template concentration^51^. CD4^+^ T cells were enriched from total PBMCs using a CD4^+^ T Cell Isolation Kit (STEMCELL Technologies, catalog 17952) and subjected to DNA extraction using commercial kits (QIAGEN DNeasy, 69504). We amplified total HIV-1 DNA using ddPCR (Bio-Rad), with primers and probes previously described (127-bp 5′-LTR-gag amplicon; HXB2 coordinates 684-810^43,52,53^ and a modified version of the intact proviral DNA assay (IPDA)^51^we published previously^54^. The droplets were subsequently read by a QX600 droplet reader, and data were analyzed using Quanta-Soft software (BIO-RAD).

### Multiple Displacement Amplification

Genomic DNA was isolated using commercial kits (QIAGEN DNeasy Blood & Tissue Kit, catalog 69504), according to the manufacturer’s instructions. Then, the genomic DNA was amplified using multiple displacement amplification (MDA) with phi29 polymerase (QIAGEN REPLI-g Single Cell Kit, catalog #150345). Buffers DLB, D1 (denaturation buffer), and N1 (neutralization buffer) were prepared according to the manufacturer’s protocol. The genomic DNA (2.5 μL) was denatured by the addition of 2.5 μL of buffer D1, followed by an incubation at room temperature for 3 mins. The mix was neutralized by adding 5 μL of buffer N1. 40 μL of the MDA master mix, consisting of nuclease-free water, REPLI-g single cell Reaction Buffer, and REPLI-g single cell DNA Polymerase, was added to the 10 μL denatured genomic DNA sample. The final 50 μL reaction was incubated at 30°C for 4 h, followed by heat inactivation at 65°C for 3 mins. The amplified product was purified using AMPure XP beads (Beckman Coulter) and was used for downstream integration site analysis.

### Enzymatic digestion of genomic DNA

This protocol was adapted and modified from the Clontech Lenti-X Integration Site Analysis Kit (catalog #631263), which, in its standard form without MDA, lacks sufficient sensitivity to detect single-copy proviral DNA in a sample. Purified MDA products were digested using three blunt-end restriction enzymes - HpaI (NEB #R0105L), SspI (NEB #R3132L), and DraI (NEB #R0129L) - to fragment genomic DNA and facilitate subsequent ligation-based integration site analysis. Briefly, 5 µL of purified MDA product was added to a master mix containing nuclease-free water, 10× restriction enzyme buffer, and either DraI or SspI (20 U µL⁻¹) or HpaI (5 U µL⁻¹), to a final reaction volume of 50 µL. Reactions were incubated at 37 °C for 18 hours to ensure complete digestion. The digested genomic DNA was then purified using AMPure XP beads and carried forward to adaptor ligation.

### Ligation of the genome-walker adaptor to the digested genomic DNA

The genomic DNA fragments are ligated with double-stranded T-linker DNA (genome walker) (see section below for exact sequence), in which the shorter strand is an oligonucleotide with a 5’ end phosphorylated (to enable efficient linker ligation) and the 3’ end is modified with an amino modification, which limits the extension of the short strand during PCR in case of linker self-ligation. Digested genomic DNA (4.8 µL) was ligated to adaptors in an 8 µL reaction containing 1.9 µL adaptor (25 µM), 0.8 µL 10× ligation buffer, and 0.5 µL T4 DNA ligase (6 U µL⁻¹; NEB #M0202L). Ligation was performed at 16 °C overnight in a thermal cycler to ensure temperature stability. The reaction was terminated by heat inactivation at 70 °C for 5 min, followed by dilution with 32 µL of DEPC-treated water (Ambion #4387937) to a final volume of 40 µL.

### Viral-host junction PCR amplification

The adaptor ligated genomic DNA fragments were used as a template for nested PCR amplification of viral-host junctions. The 5′LTR HIV-1 junction was amplified using LSP1 (GCTTCAGCAAGCCGAGTCCTGCGTCGAG) and LSP2 (GCTCCTCTGGTTTCCCTTTCGCTTTCAA) as forward primers, both derived directly from the CloneTech LentiX kit. The 3′LTR HIV-1 junction was amplified using LestralLTR1 (CTTAAGCCTCAATAAAGCTTGCCTTGAG) and LestralLTR2 (AGACCCTTTTAGTCAGTGTGGAAAATC)^5^. These were paired with adaptor-specific reverse primers AP1 (GTAATACGACTCACTATAGGGC) and AP2 (ACTATAGGGCACGCGTGGT) for the first and second PCR reactions, respectively. The AP1 primer is specific to the 5’ single-stranded region of the T-linker to avoid unwanted amplification of the host DNA. The adaptor sequence was adapted from the CloneTech LentiX kit (catalog #631263).

Each 25 µL PCR reaction contained 1× reaction buffer, 1× dNTP mix, 0.2 µM of each primer, and Advantage 2 polymerase (Clontech Advantage® 2 PCR Kit, Cat. #639206). For the first PCR, 15 µL of master mix was combined with 10 µL of diluted adaptor-ligated genomic DNA, using AP1/LSP1 for 5′LTR junctions and AP1/LestralLTR1^5^ for 3′LTR junctions. Cycling conditions were: 7 cycles of 94 °C for 25 s and 72 °C for 3 min, followed by 32 cycles of 94 °C for 25 s and 67 °C for 3 min, and a final extension at 67 °C for 7 min.

For the second (nested) PCR, 24 µL of master mix was combined with 1 µL of first-round PCR product, using AP2/LSP2 for 5′LTR junctions and AP2/LestralLTR2^5^ for 3′LTR junctions. Cycling conditions were: 5 cycles of 94 °C for 25 s and 72 °C for 3 min, followed by 20 cycles of 94 °C for 25 s and 67 °C for 3 min, with a final extension at 67 °C for 7 min. Both 5′LTR and 3′LTR viral-host junctions were successfully amplified under these conditions. The resulting PCR products were subjected to Illumina MiSeq paired-end 150 base pairs (bp) sequencing.

### Cell culture

Jurkat cells and MOLT-4 cells were cultured at 37 °C and 5% CO2 in RPMI supplemented with 10 % FCS and Gentamycin. All cell lines were monitored regularly by the SG-PERT assay^55^ to ensure the absence of retroviral contamination. Cell lines used in this study were devoid of mycoplasma contamination.

## Availability

BulkIntSiteR is available under an open-source license at: https://github.com/guineverelee/BulkIntSiteR/

## Acknowledgements

We acknowledge the Massachusetts General Hospital Center for Computational and Integrative Biology DNA Core for Illumina sequencing support. This work was supported by NIH grants Research Enterprise to Advance a Cure for HIV (REACH) UM1AI164565 (RBJ), R21AI150398 (GQL), R01AI162221 (GQL), and the Center for the Structural Biology of HIV RNA U54 AI170660 (PB). We acknowledge Dr. Douglas Barrows, Bioinformatics Analyst, Bioinformatics Resource Center, The Rockefeller University, New York. The authors gratefully acknowledge the MGH DNA core facility, Massachusetts, USA. This article is subject to HHMI’s Open Access to Publications policy. HHMI lab heads have previously granted a nonexclusive CC BY 4.0 license to the public and a sublicensable license to HHMI in their research articles. Pursuant to those licenses, the author-accepted manuscript of this article can be made freely available under a CC BY 4.0 license immediately upon publication.

---

**Figure S1.**
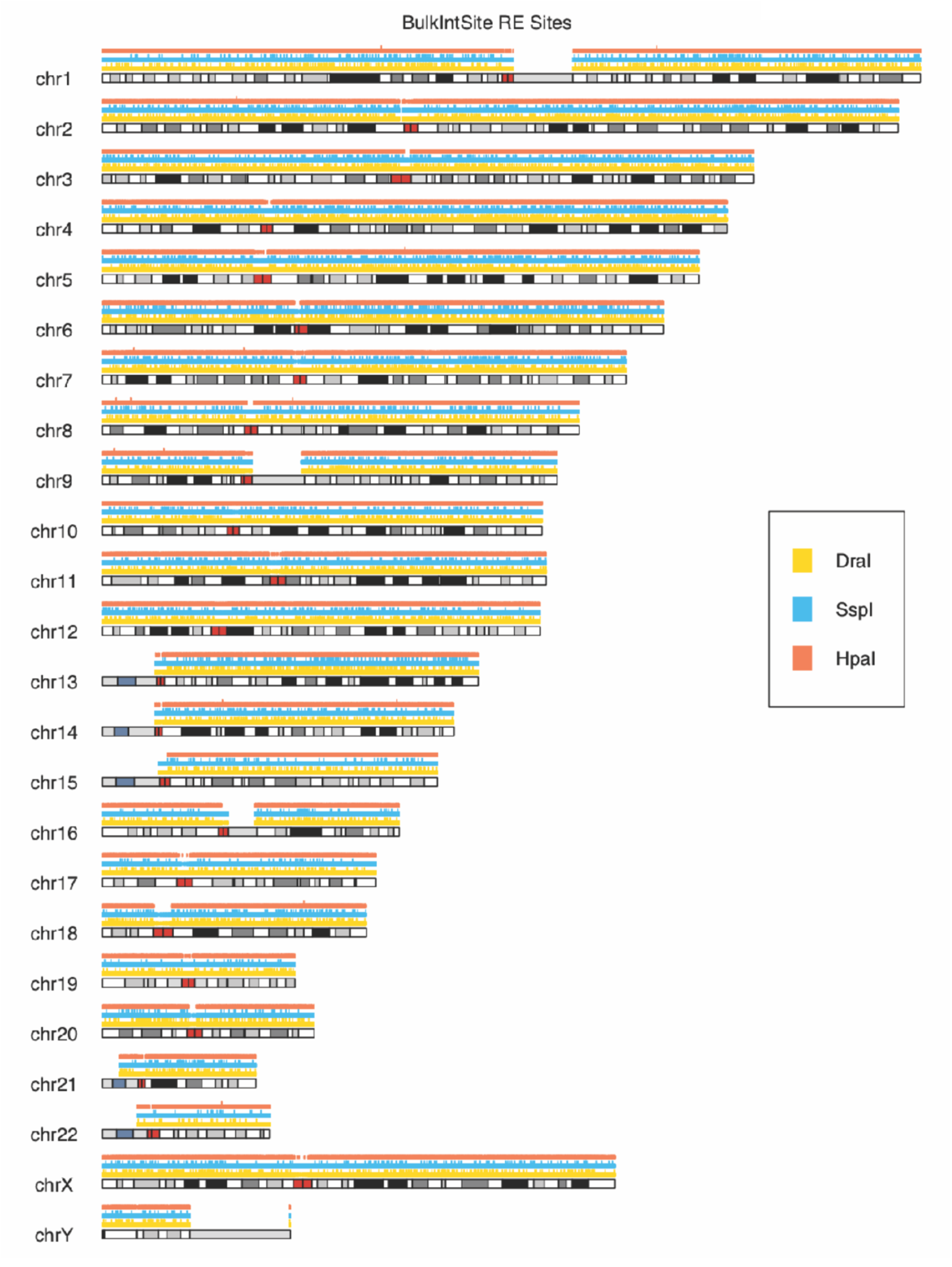
Genome-wide distribution of the restriction enzyme recognition site. The karyoplot shows the DraI (yellow), SspI (cyan), and HpaI (orange) restriction sites distribution genome-wide on the human reference genome assembly hg38. *RE, restriction enzymes*.

**Figure S2.**
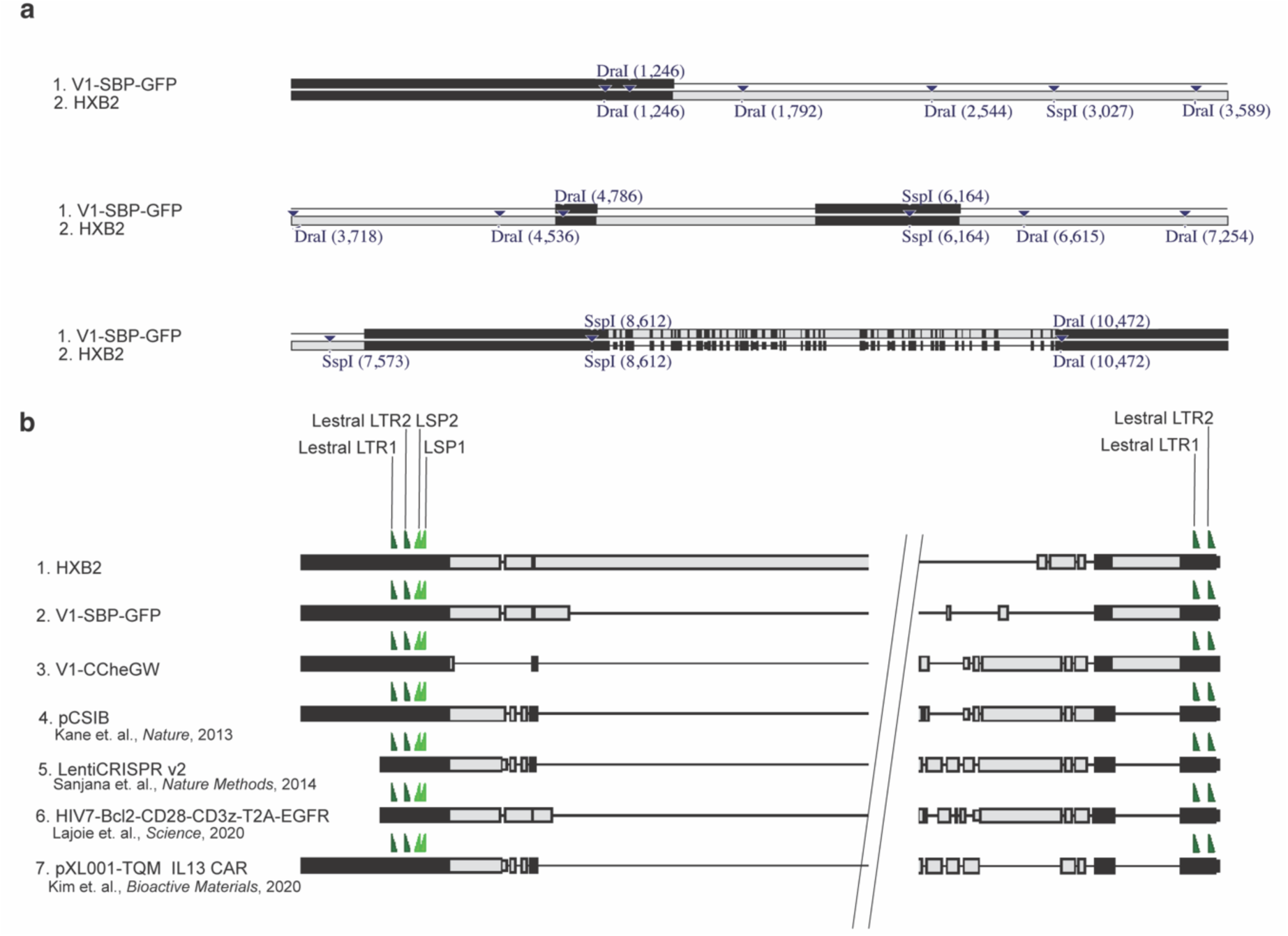
Distribution of restriction enzyme sites within the provirus genome and PRISM-seq primer binding sites compatibility across HIV and lentiviral vector delivery systems. **(a)** Location of DraI and SspI restriction sites in the genome of the full-length HIV-1 reference sequence (HXB2) and replication-deficient HIV-1 reporter (V1-SBP-GFP) from 5′ to 3′ direction. Restriction sites coordinates in the proviral genome are mentioned within brackets next to each restriction enzyme. Black and gray colors represent the identical and non-identical sequences, respectively, between the V1-SBP-GFP and HXB2 genome sequence. **(b)** Multiple sequence alignment of HXB2, replication-deficient single-round HIV-1 reporters (V1- SBP-GFP and V1-CCheGW), pCSIB (lentivirus-based overexpression construct to generate stable cell lines), LentiCRISPRv2 (CRISPR/Cas9-based genome editing lentiviral vector), HIV7-Bcl2-CD28-CD3z-T2A-EGFR, and pXL001-TQM IL13 CAR lentiviral vectors. Primer binding sites for 5′LTR (LSP1 and LSP2) and 3′LTR (LestralLTR1 and LestralLTR2) viral-host junction amplification are conserved across these sequences.

**Figure S3.**
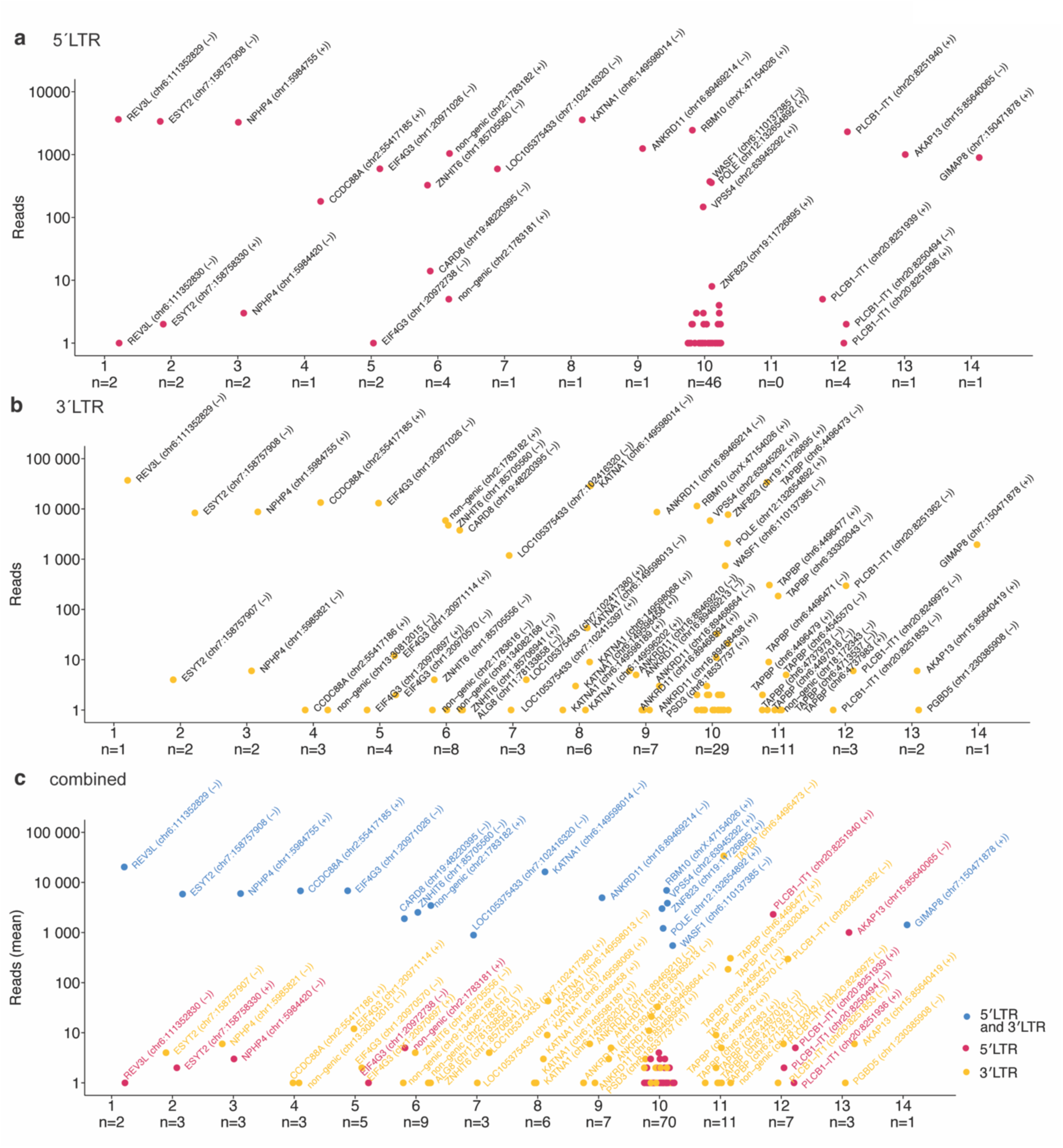
Identification of proviral integration sites using PRISM-seq in putative single-cell clones #1-14, raw pre-filter data. **(a-c)** Proviral integration sites retrieved from individual putative clones, either from the 5′LTR (a) or 3′LTR (b) or combined (c) viral host junction. Blue, red, and yellow dots represent the integration sites identified by both 5′LTR and 3′LTR, exclusively by 5′LTR or 3′LTR, respectively. LTR; long terminal repeat of lentiviruses.

**Figure S4.**
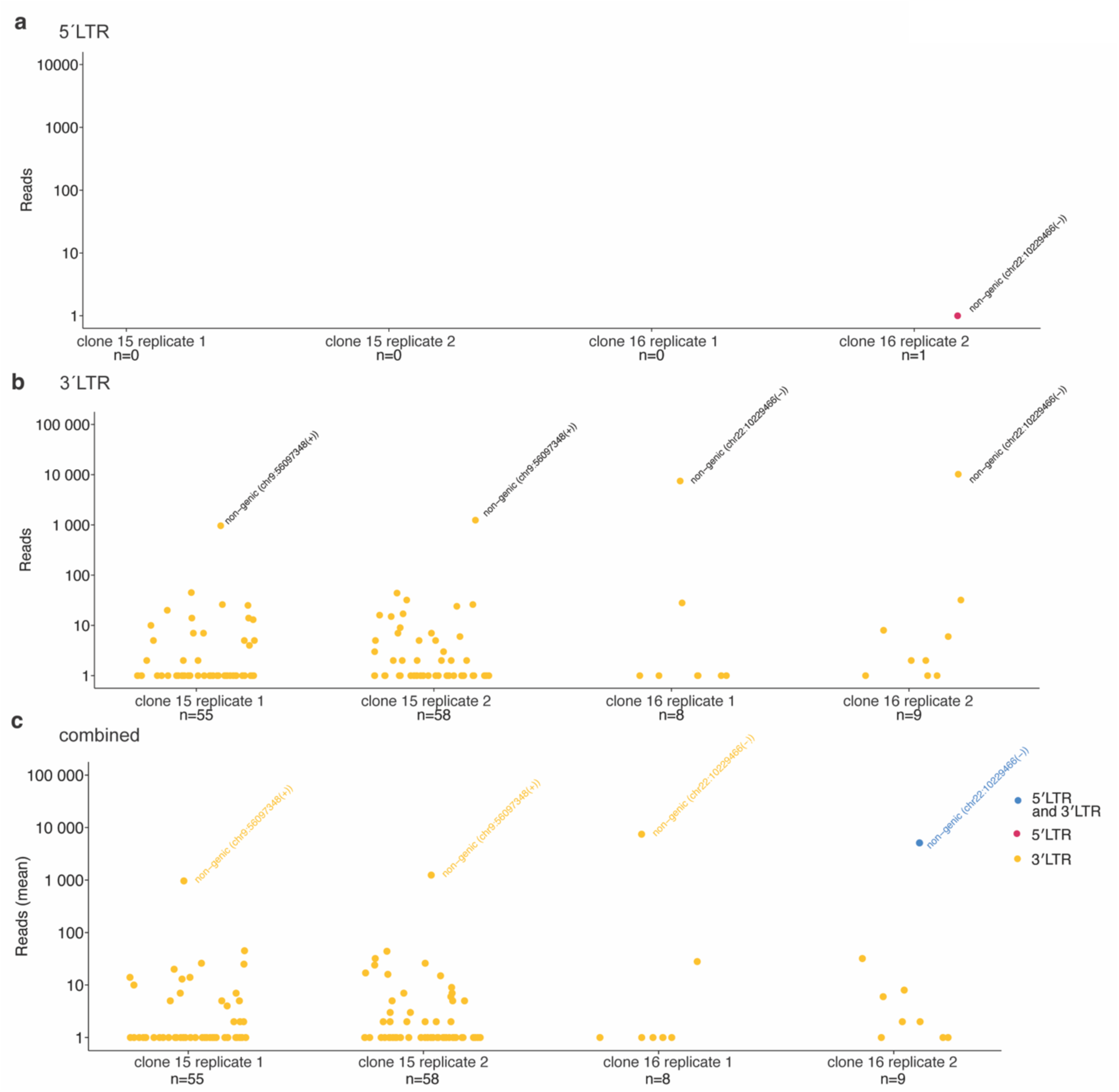
Identification of proviral integration sites using PRISM-seq in clones with known centromeric integrations #15 and #16, raw pre-filter data. (a-c) Proviral integration sites were identified in clones #15 and #16, either from the 5′LTR (a) or 3′LTR (b) or combined (c) viral host junction using PRISM-seq. Results for clones #15 and #16 are shown for both technical duplicate reactions.

**Figure S5.**
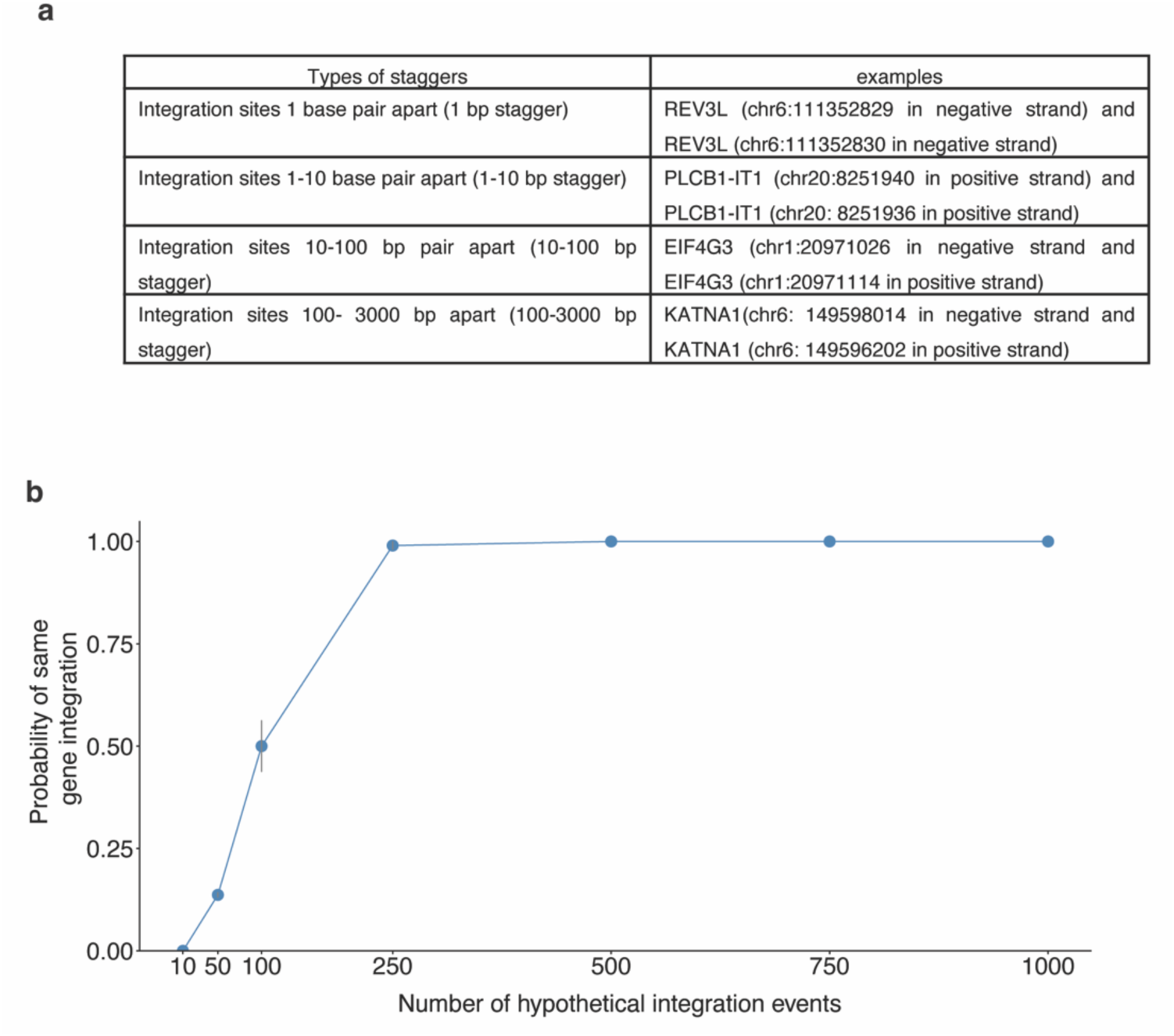
Identification of look-alike integration site artifact cases and the relationship between integration events and the probability of having true multiple integrations into the same gene. **(a)** Characterization of artifacts generated during the integration sites assay. Each staggered case is distinguished based on its genomic distance (bp) from the high-confidence integration sites coordinates of a putative clone. The examples shown here were observed in distinct putative clones. **(b)** Regarding Step 3 in our proposed quality filter, which involves collapsing integration sites that map to the same gene when found within a single reaction: We recognize that when PRISM-seq is applied to non-clonal clinical samples, there is a genuine possibility that proviruses have integrated into distinct locations within the same gene. The likelihood of detecting genuine same gene integration depends on the number of integration events sampled within a PRISM-seq reaction. We performed a Monte Carlo simulation and confirmed that the likelihood of detecting genuine same-gene integration was 0%, 0%, 14%, 50%, and 99% when 1, 10, 50, 100, and 250 unique integration events were sampled. Therefore, we suggest that users should quantify targeting viral DNA copies by techniques such as ddPCR and turn off Step 3 if more than 50 target copies were input into each PRISM-seq reaction, with the caveat that specificity would be compromised. In this study, all reactions were performed at 50 copies per reaction input. *Bp, base pairs*.

**Figure S6.**
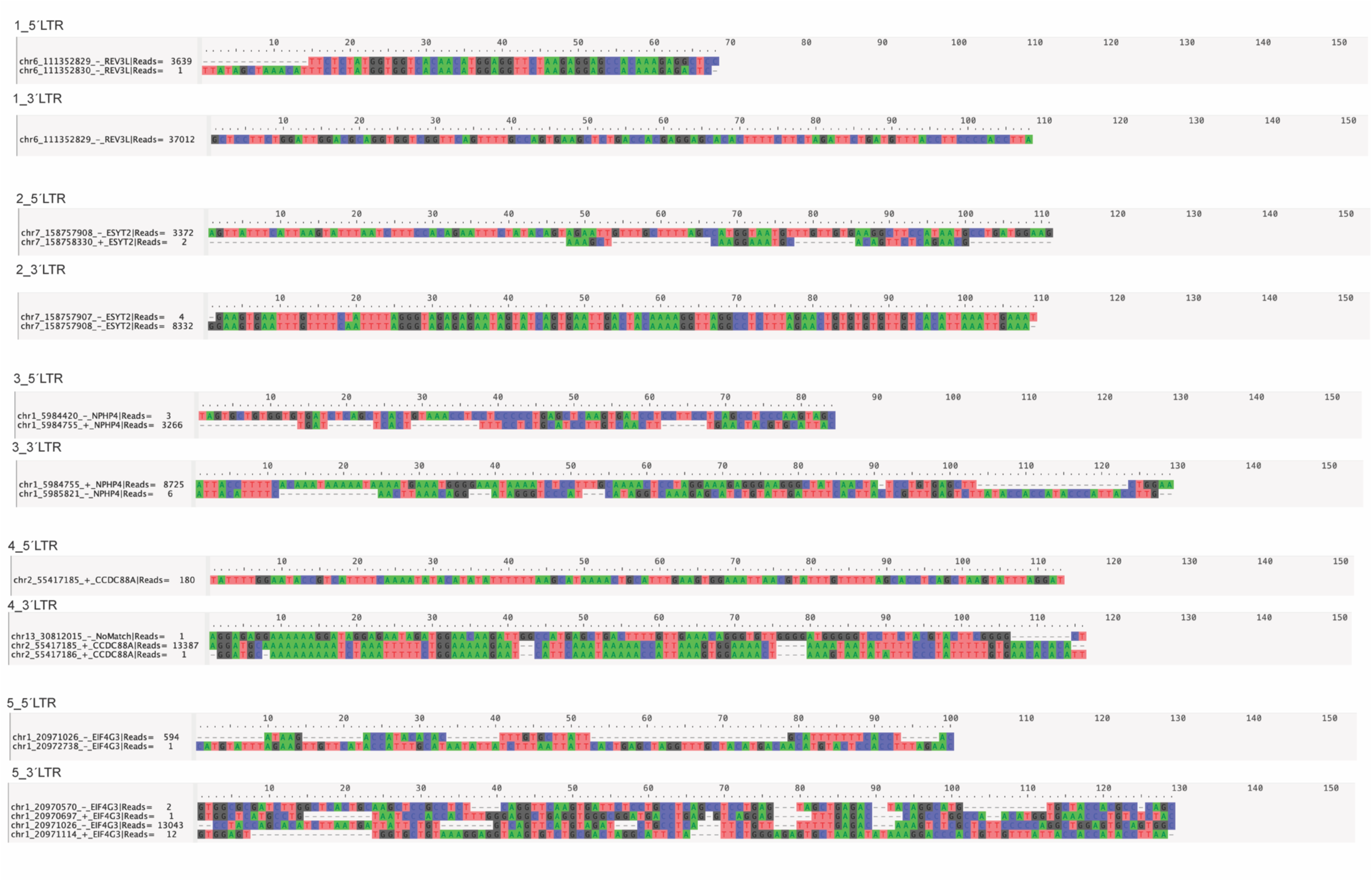
Comparison of host sequences for each unique integration site identified within the putative clones. Low-confidence integration sites shared the same junction sequence as the corresponding high-confidence integration site in clones #1, #2, and #4.

**Figure S7.**
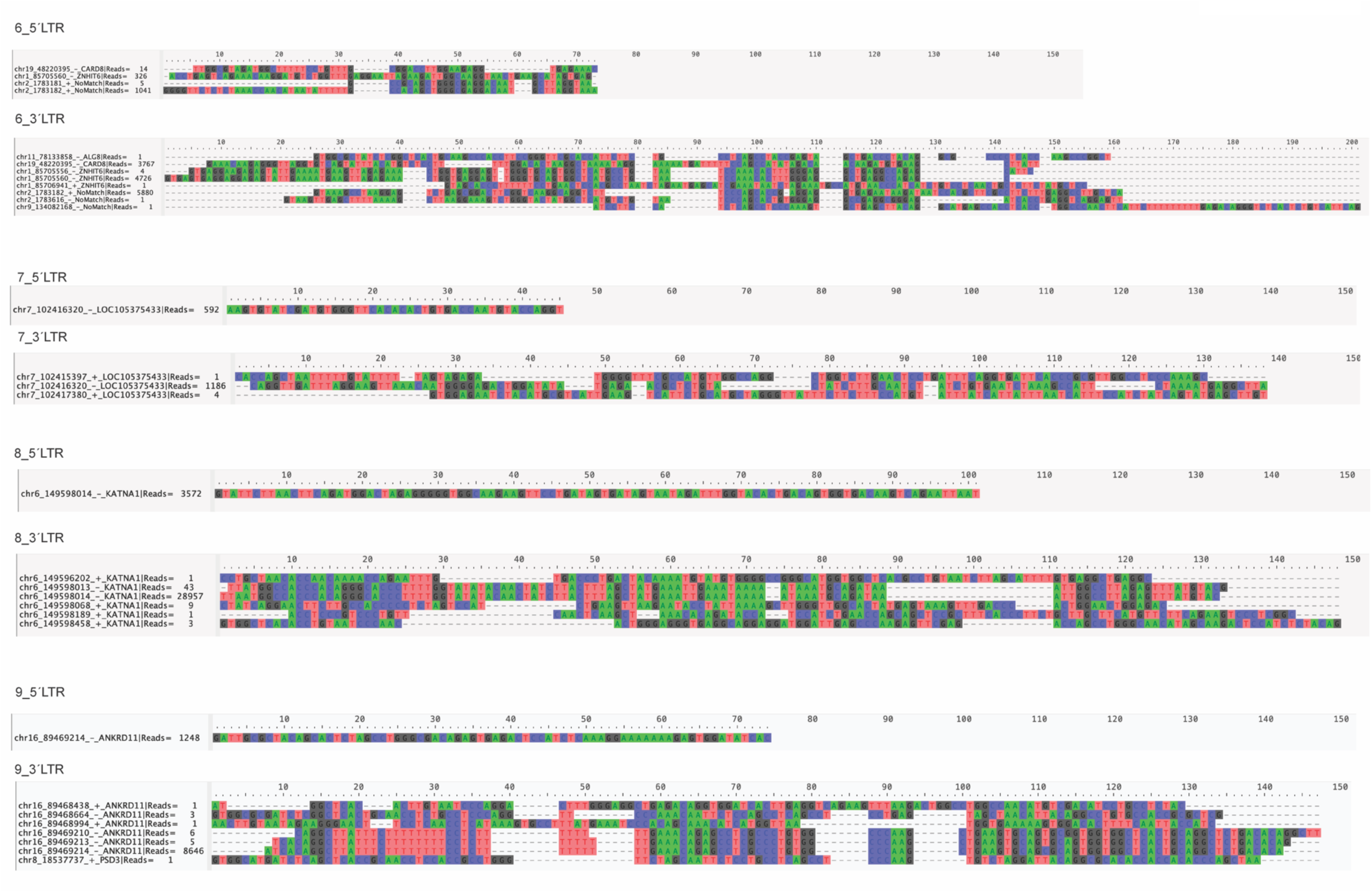
Comparison of host sequences for each unique integration site identified within the putative clones. Low-confidence integration sites shared the same junction sequence as the corresponding high-confidence integration site in clones #6, #8, and #9.

**Figure S8.**
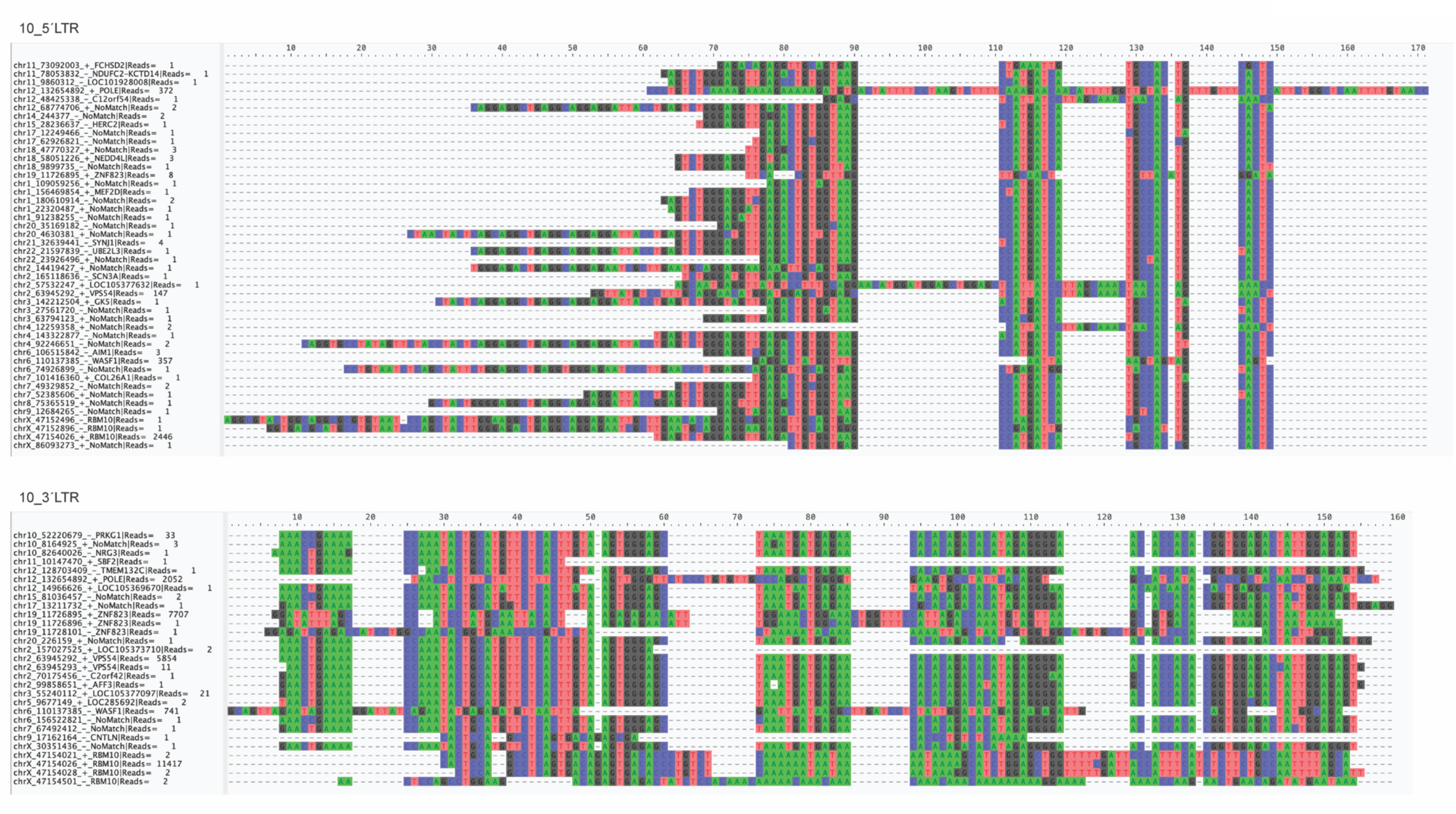
Comparison of host sequences for each unique integration site identified within the putative clones. Low-confidence integration sites shared the same junction sequence as the corresponding high-confidence integration site in clone #10.

**Figure S9.**
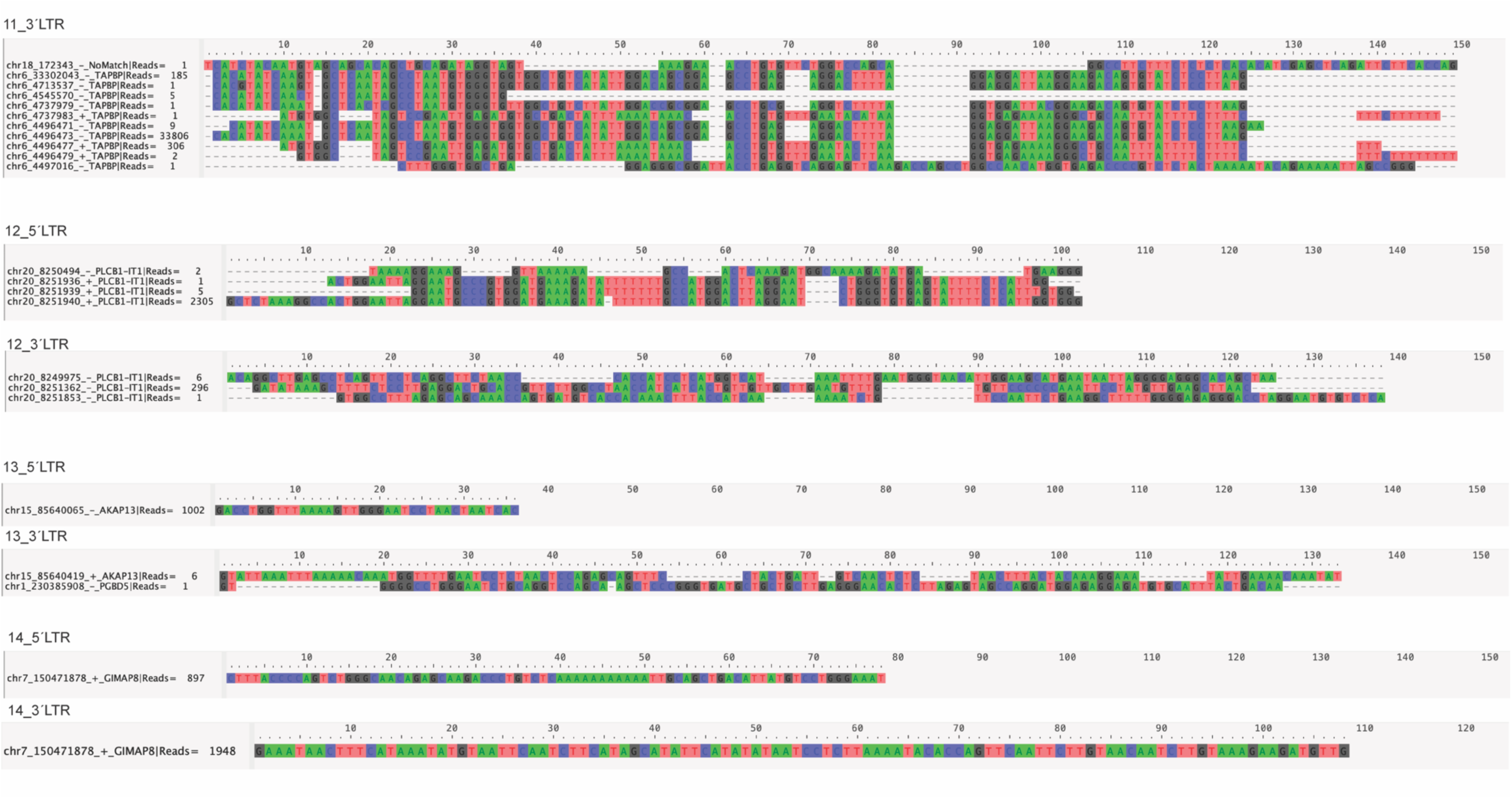
Comparison of host sequences for each unique integration site identified within the putative clones. Low-confidence integration sites shared the same junction sequence as the corresponding high-confidence integration site in clones #11 and #12.

**Figure S10.**
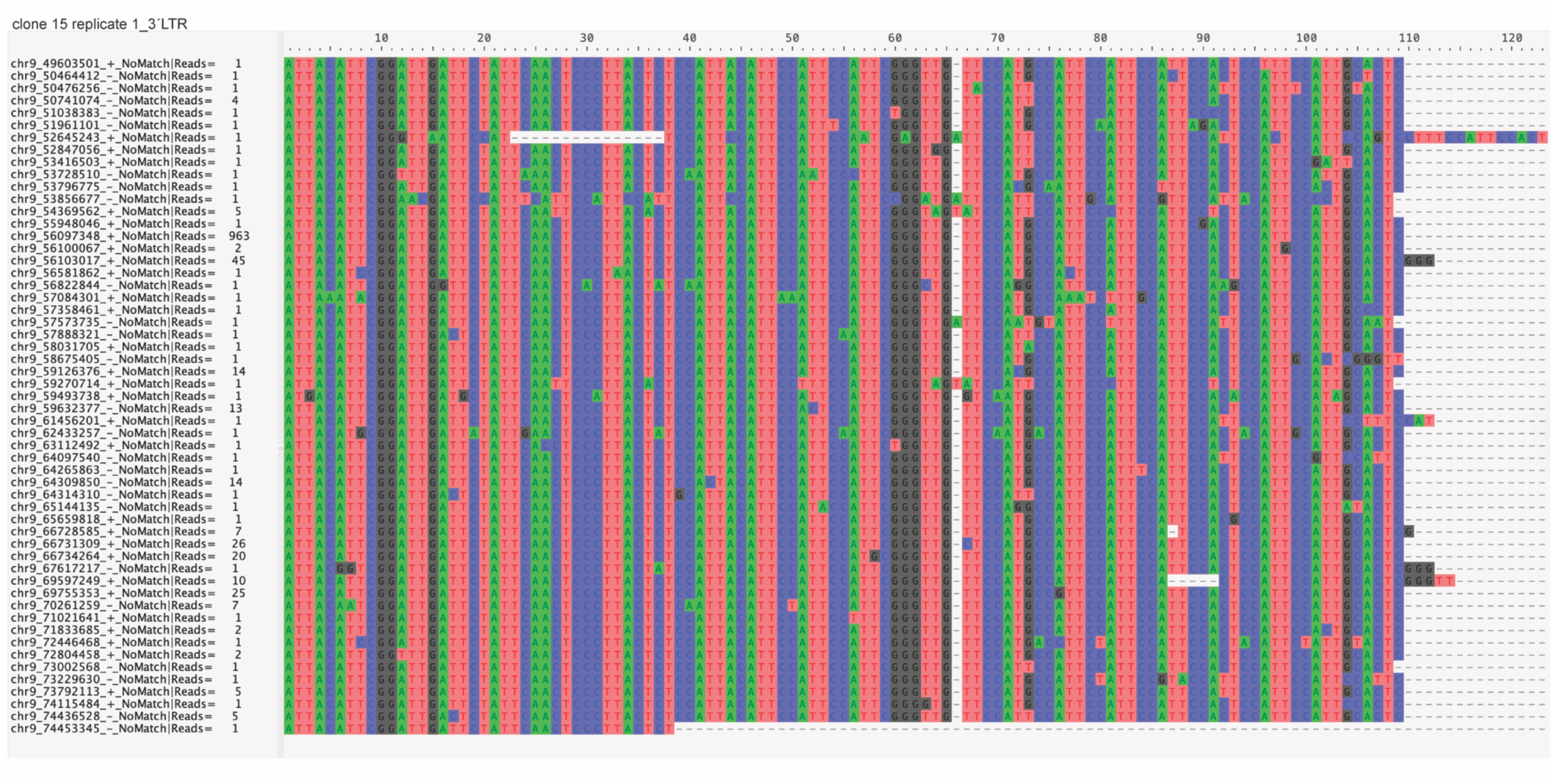
Comparison of host sequences for each unique integration site identified in a clone associated with centromeric integration. Low-confidence integration sites shared the same junction sequence as the corresponding high-confidence integration site in clone #15, replicate 1.

**Figure S11.**
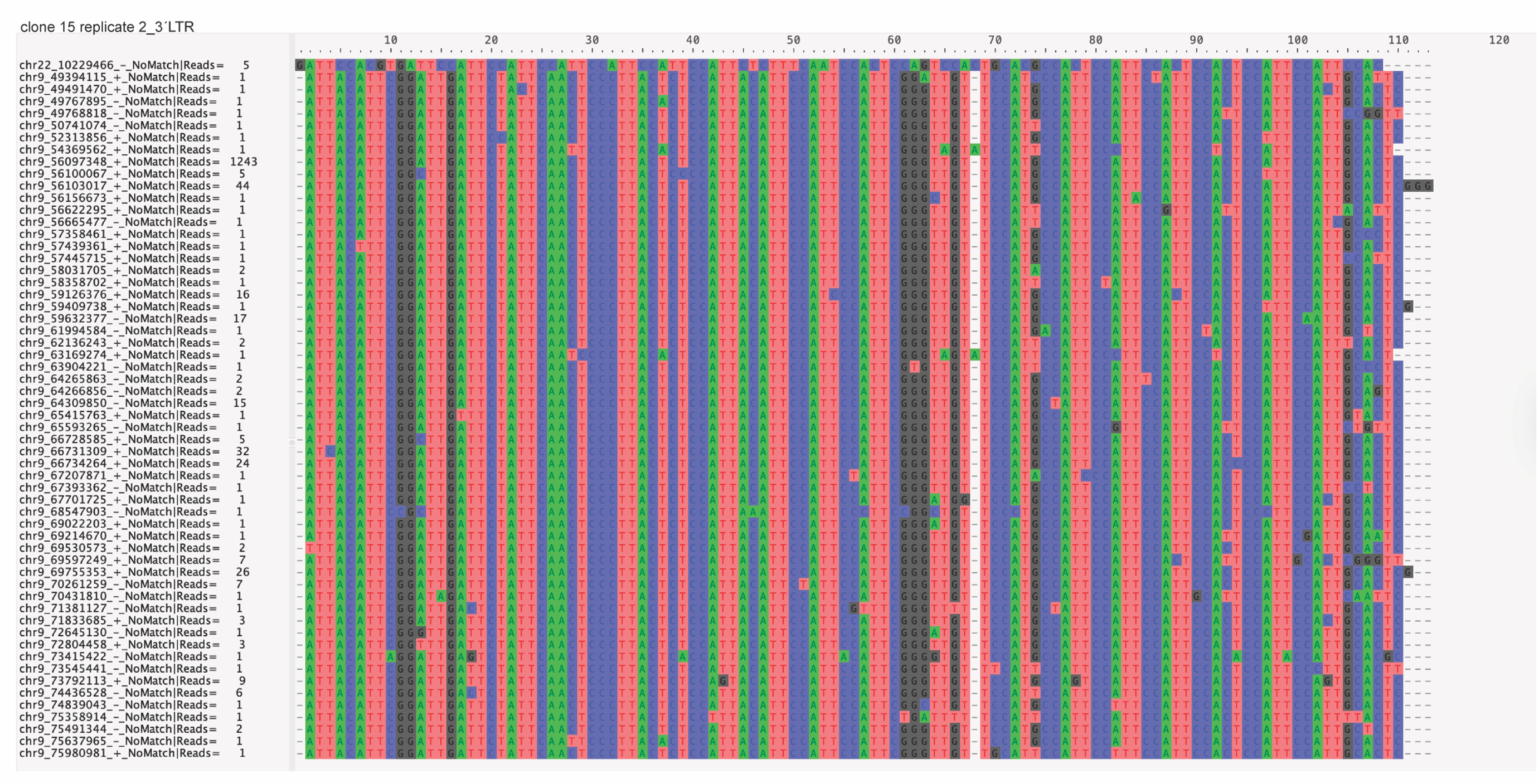
Comparison of host sequences for each unique integration site identified in a clone associated with centromeric integration. Low-confidence integration sites shared the same junction sequence as the corresponding high-confidence integration site in clone #15, replicate 2.

**Figure S12.**
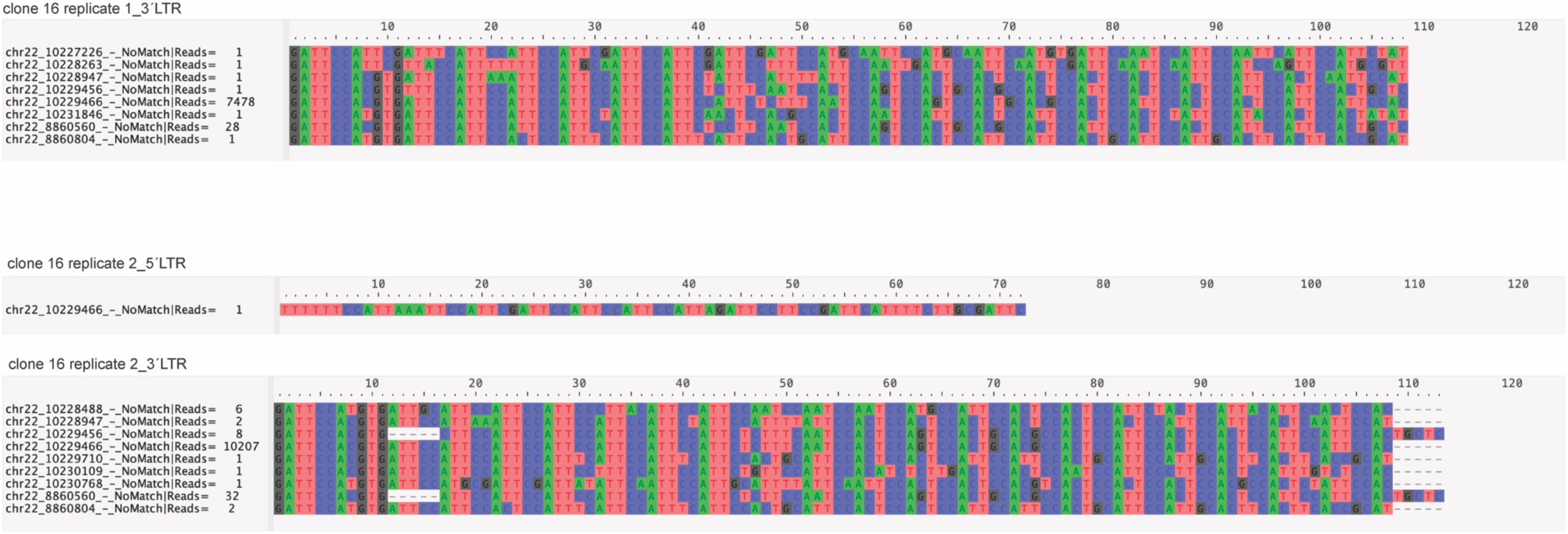
Comparison of host sequences for each unique integration site identified in a clone associated with centromeric integration. Low-confidence integration sites shared the same junction sequence as the corresponding high-confidence integration site in clone #16.

**Figure S13.**
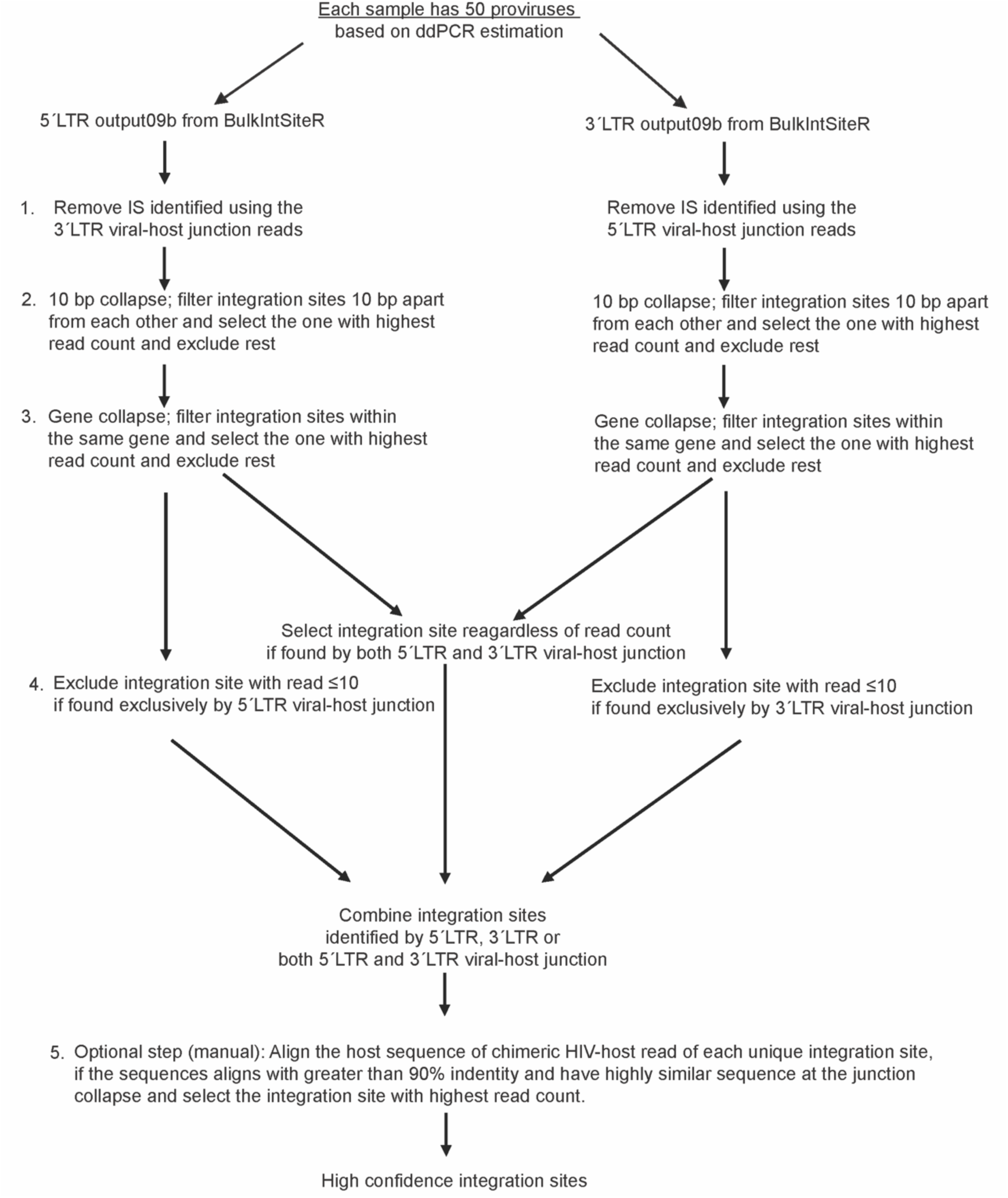
Data-guided noise removal approach. Leveraging the clonal nature of Clones #1-16, we devised a quality control filter to remove assay noise. This set of quality filters is applied to the BulkIntSiteR output; (1) filter and remove the spillover noise of integration site identified by 3′ LTR in the 5′ LTR library and vice versa. (2) integration sites within ≤10 bp of each other were collapsed by retaining the site with the highest read count; (3) integration sites located within the same gene were filtered to retain only one site with the highest read count, excluding all others if the input template count is ≤50 copies; (4) integration sites detected exclusively by either 5′ LTR or 3′ LTR junctions were excluded if supported by ≤10 read counts. Integration sites detected by both 5′ LTR and 3′ LTR virus-host junctions, regardless of the raw read counts, and the sites that passed step 4 were classified as high-confidence. An optional Step (5), involving user-guided homology clustering of host sequences derived from the viral-host chimeric reads, removes look-alike integration sites that share identical junction sequences and co-occur within the same reaction, retaining only the site with the highest read count. Steps 1-4 are fully automated, and the accompanying script is included with the BulkIntSiteR package; Step 5 requires user input and manual curation.

**Figure S14.**
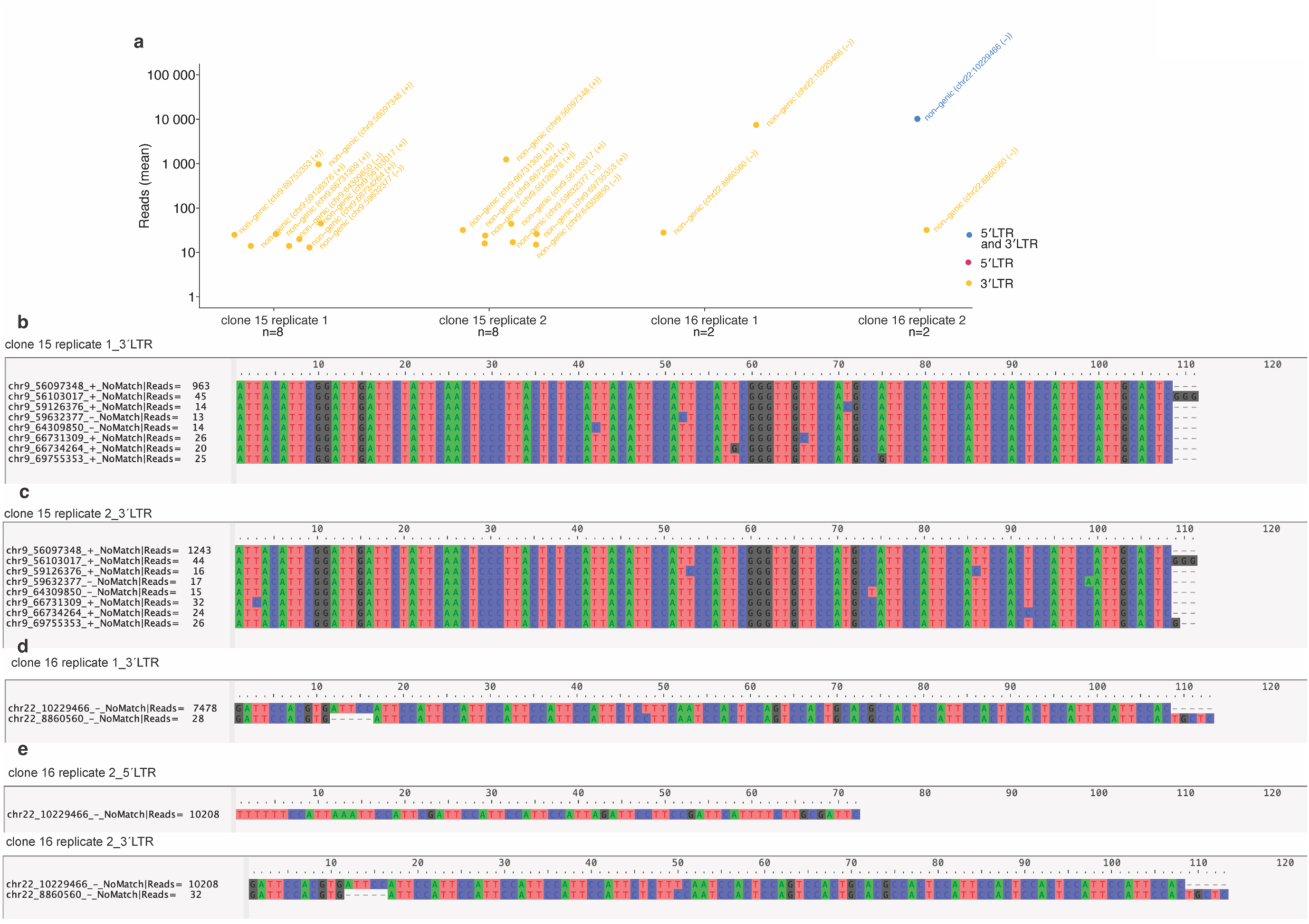
Post-filter recovery of proviral integration sites found within centromeric clones #15 and #16. **(a)** Integration sites identified by the BulkIntSiteR software were processed using the QC filtering pipeline (with steps 1-4 applied), and results for clones #15 and #16 are shown for technical duplicate reactions. **(b-e)** Comparison of the host sequence for all unique integration sites after QC filtering for clones #15 and #16. The unexpected low-confidence integration sites share the same junction sequence as the high-confidence integration site for clones #15 and #16.

**Figure S15.**
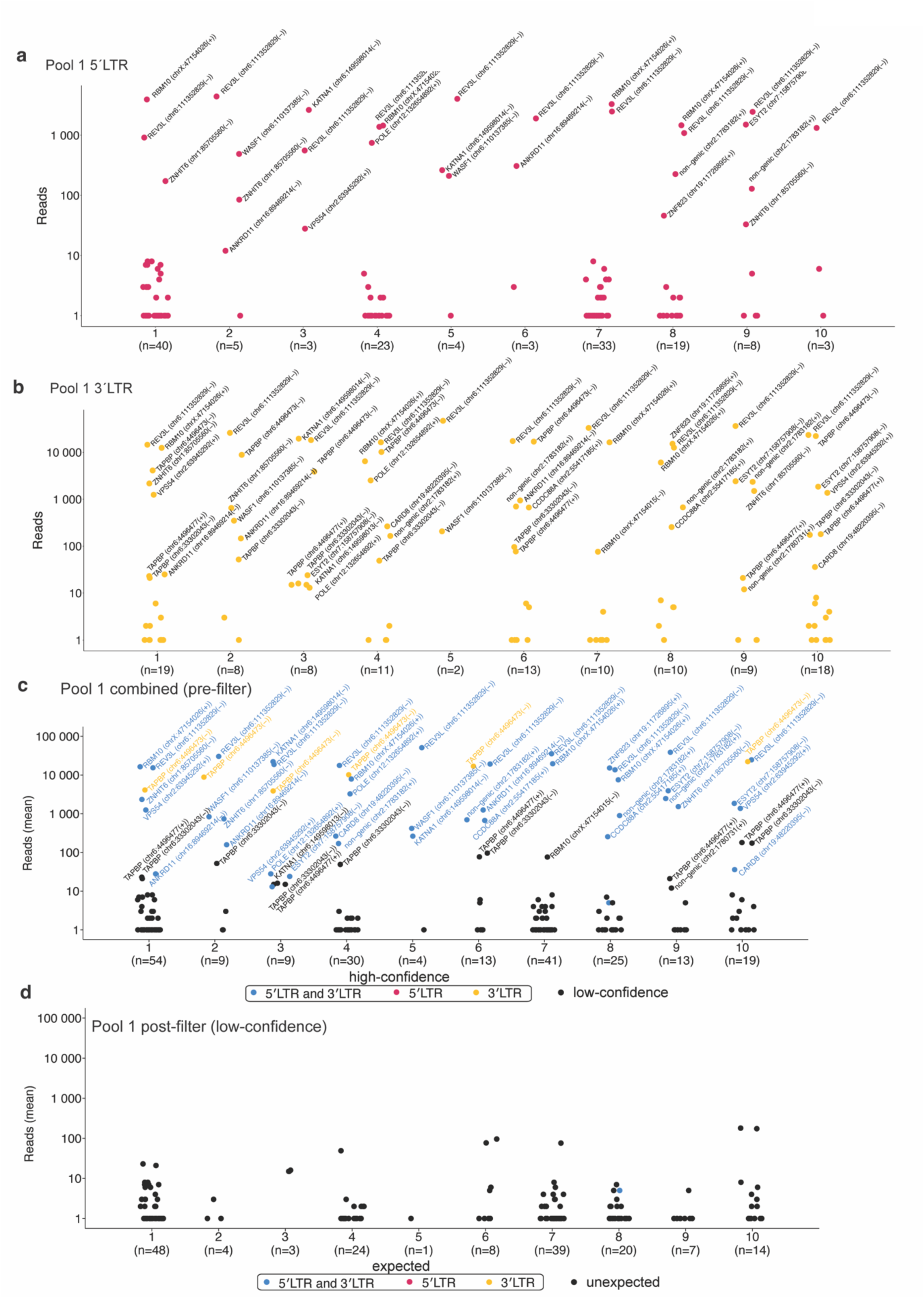
Proviral integration sites identification in pool 1. **(a-c)** Dot plots representing the total number of unique integration sites identified in each replicate of pool 1 by either the 5′LTR (a) or 3′LTR (b) or combined (c) viral-host junction. The total number of unique integration sites retrieved in an individual replicate is represented as an n value. Blue, red, and yellow dots represent integration sites identified by both 5′LTR and 3′LTR, exclusively by 5′LTR or exclusively by 3′LTR PRISM-seq reactions, respectively. (d) Post-filter identification of low-confidence expected and unexpected integration sites.

**Figure S16.**
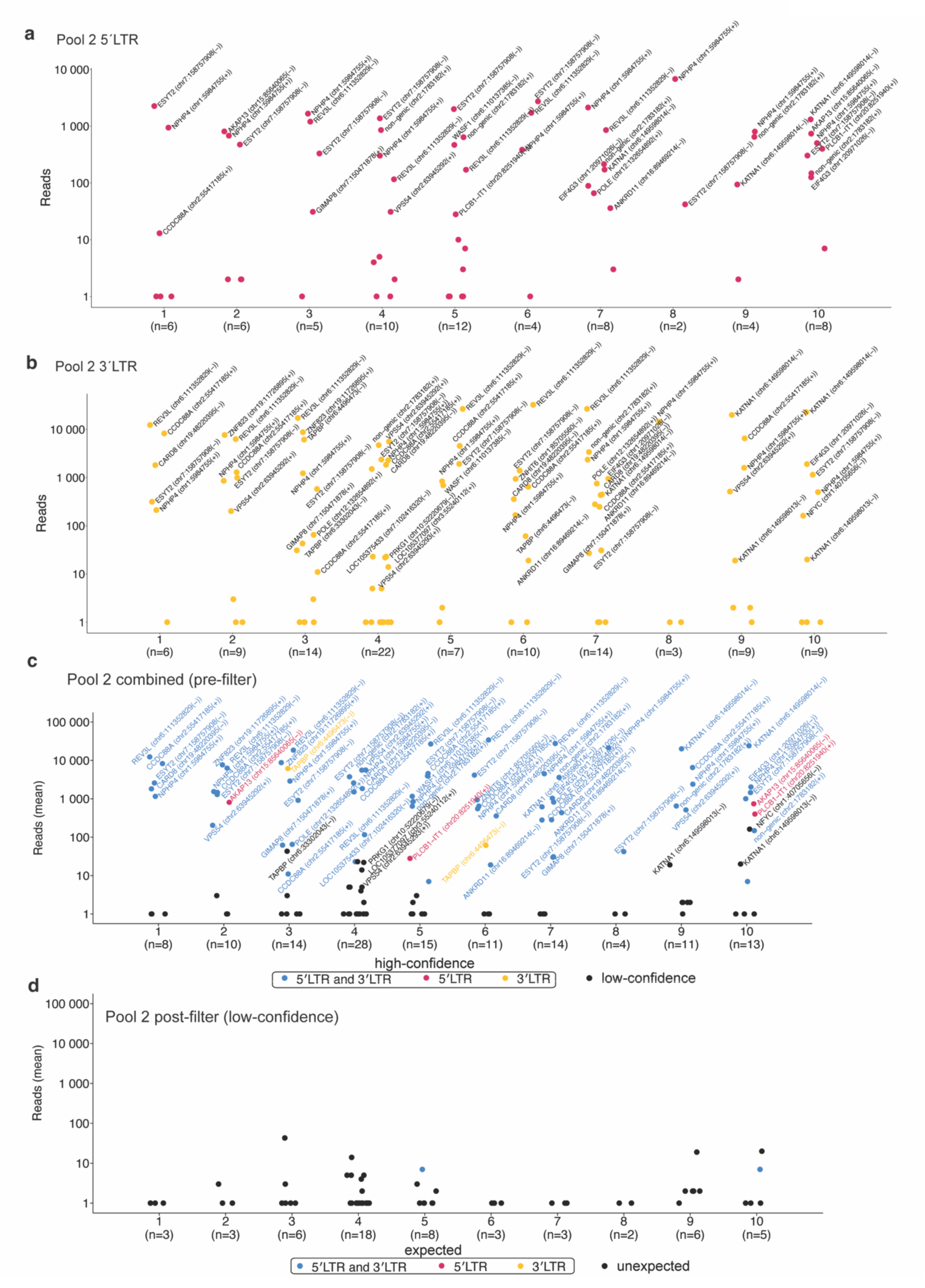
Proviral integration sites identification in pool 2. **(a-c)** Dot plots representing the total number of unique integration sites identified in each replicate of pool 2 by either the 5′LTR (a) or 3′LTR (b) or combined (c) viral-host junction. The total number of unique integration sites retrieved in an individual replicate is represented as an n value. Blue, red, and yellow dots represent the integration sites identified by both 5′LTR and 3′LTR, exclusively by 5′LTR or exclusively by 3′LTR PRISM-seq reactions, respectively. (d) Post-filter identification of low-confidence expected and unexpected integration sites.

**Figure S17.**
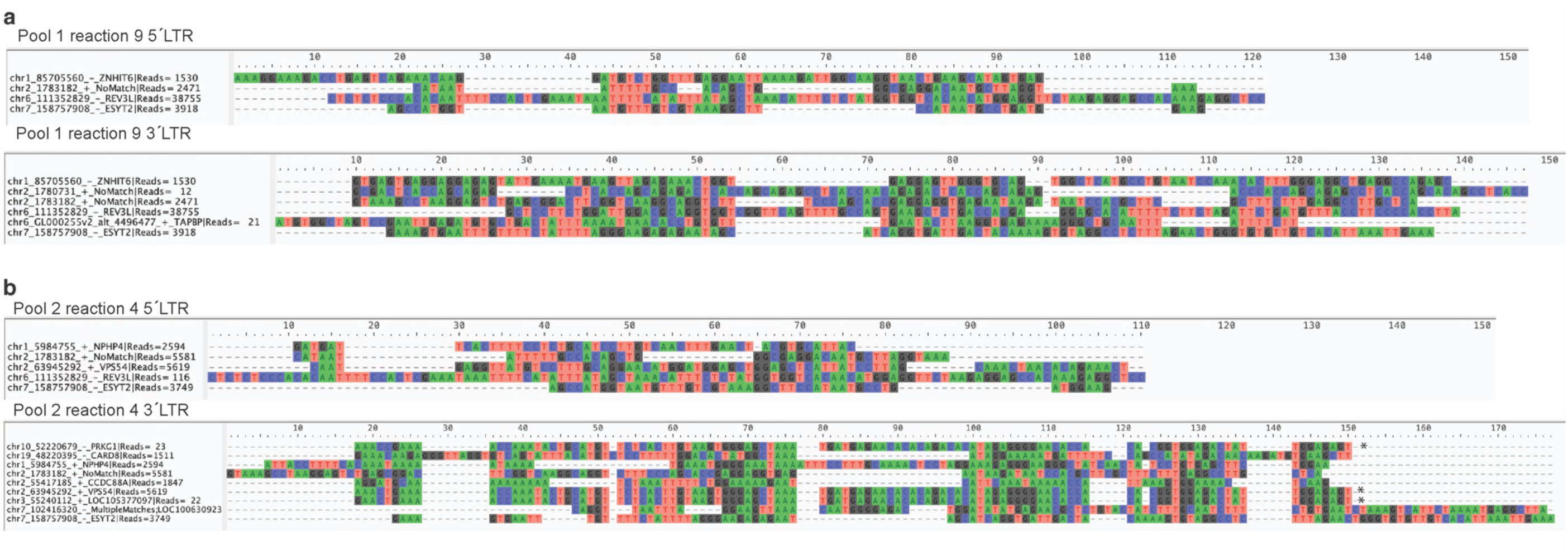
Comparison of host sequences of unexpected integration sites observed in pool 1 and pool 2, respectively (a and b). Host sequence derived from unexpected integration sites found in pool 1 replicate 9 (a) and pool 2 replicate 4 (b) by both 5′LTR and 3′LTR PRISM-seq reactions. Unexpected integration sites sharing the same junction sequence as the high-confidence site and/or are ±10 kb away from the expected high-confidence integration site are labelled with an asterisk (*).

